# Single cell compendium of muscle microenvironment in peripheral artery disease reveals altered endothelial diversity and LYVE1^+^ Macrophage Activation

**DOI:** 10.1101/2023.06.21.545899

**Authors:** Guillermo Turiel, Thibaut Desgeorges, Evi Masschelein, Zheng Fan, David Lussi, Christophe M. Capelle, Giulia Bernardini, Raphaela Ardicoglu, Katharina Schönberger, Manuela Birrer, Sandro F. Fucentese, Jing Zhang, Daniela Latorre, Stephan Engelberger, Katrien De Bock

## Abstract

Peripheral artery disease (PAD) results from atherosclerosis and chronic narrowing of lower limb arteries leading to decreased muscle perfusion. Current treatments are suboptimal, partly due to limited understanding of PAD muscle pathology. Here we use scRNAseq and spatial transcriptomics to analyze the composition of the muscle microenvironment in non-ischemic and PAD patients. We identified *ATF3/4^+^* endothelial cells (ECs) that exhibit altered angiogenic and immune regulatory profiles during PAD and confirmed that ATF4 signaling in ECs is required for effective ischemia recovery. Also, capillary ECs display features of endothelial-to-mesenchymal transition. Furthermore, *LYVE1*^hi^MHCII^low^ macrophages are the dominant macrophage population in human muscle, adopting a more pro-inflammatory profile during PAD. Finally, we analyzed alterations in intercellular communication within the muscle microenvironment during PAD and confirmed that EC-derived factors can influence macrophage polarization. This dataset deeply characterizes the PAD muscle microenvironment and provides a resource for exploration of targeted therapies.

## Introduction

Peripheral artery disease (PAD) is caused by atherosclerosis and chronic narrowing of lower limb arteries leading to decreased muscle perfusion and oxygenation. Cardiovascular risk factors, such as hypertension, hyperlipidemia and diabetes mellitus are important driving risk factors of PAD and patients often present other cardiovascular diseases^1^. PAD patients suffer from pain and fatigue in the limb muscles due to exercise-induced ischemia which resolves after a short period of rest, a syndrome termed intermittent claudication (IC)^2,3^. If left untreated or in more severe stages, persistent lack of limb perfusion can lead to critical limb ischemia (CLI), which comprises rest pain, tissue loss and necrosis, and ulcerations, which often progresses to amputation^1^. Current guidelines for treating PAD target large atherosclerotic lesions in the feeding arteries, but long-term clinical outcomes have been suboptimal, and many patients suffer from adverse events^4^.

One potential reason for this might be that there is underlying skeletal muscle and vascular pathology in PAD^3^. Indeed, microvascular dysfunction has been reported, with abnormal microvascular architecture and fibrosis and few studies recently reported partial endothelial-to-mesenchymal (EndoMT) transition in PAD patients^5,6^ and capillaries of end-stage patients^6^. Also, impaired vasoreactive responses might impair oxygen and nutrient delivery into the ischemic muscle^7^. Interestingly, some^8–11^ but not all^12,13^ studies observed increased vascular density. While the discrepancy underlying these conflicting observations is unclear, persistent ischemia in PAD patients despite increased vascularization nonetheless suggests that an active angiogenic response does not resolve tissue ischemia.

Studies using preclinical muscle injury models showed that microvascular endothelial cells (ECs) are not only essential for inducing muscle revascularization, ECs also steer muscle regeneration by interacting with other cells in their microenvironment. ECs activate muscle stem cells^14^, control vascular tone through interaction with smooth muscle cells^15^ and control the entry of immune cells into the tissue, their differentiation into specialized immune effectors as well as (co-)define their functional properties^16,17^. However, our understanding on how ECs interact with other cell types and might contribute to skeletal muscle pathology in PAD, remains largely unknown.

Macrophages are key regulators of muscle regeneration^18^ but also stimulate muscle angiogenesis as well as arteriogenesis in preclinical PAD models^16,19^. The genetic and functional contribution of macrophages to skeletal muscle pathology in PAD are however unclear. Higher levels of inflammatory cytokines correlated with shorter onset claudication pain^20^, but assessments of macrophage numbers and properties in skeletal muscle showed conflicting results with one manuscript showing a majority of pro-inflammatory CD80^+^ macrophages^21^, while others reported higher CD11b^+^CD206^+^ regenerative macrophages^22^

In this study, we performed a thorough transcriptomic and functional characterization of the mononuclear cell composition in skeletal muscle during PAD. We aimed to better understand the contribution of the muscle microenvironment to PAD.

## Results

### scRNAseq reveals a high degree of cellular heterogeneity in human skeletal muscle

To investigate the cellular landscape of PAD patients, we collected gastrocnemius muscle samples from individuals undergoing either PAD bypass surgery (PAD) or co-morbidity matched non-ischemic individuals undergoing lower limb aneurysm surgery (non-ischemic) (see Methods, Fig. 1a, Table 1). Non-ischemic patients had a normal ankle-brachial index (ABI) and no clinical symptoms of PAD (no claudication symptoms, Fontaine classification = I) (Table 1). Pre-operative angiographic imaging revealed that only PAD patients presented severe popliteal occlusions (GLASS grade III).

**Fig. 1:**
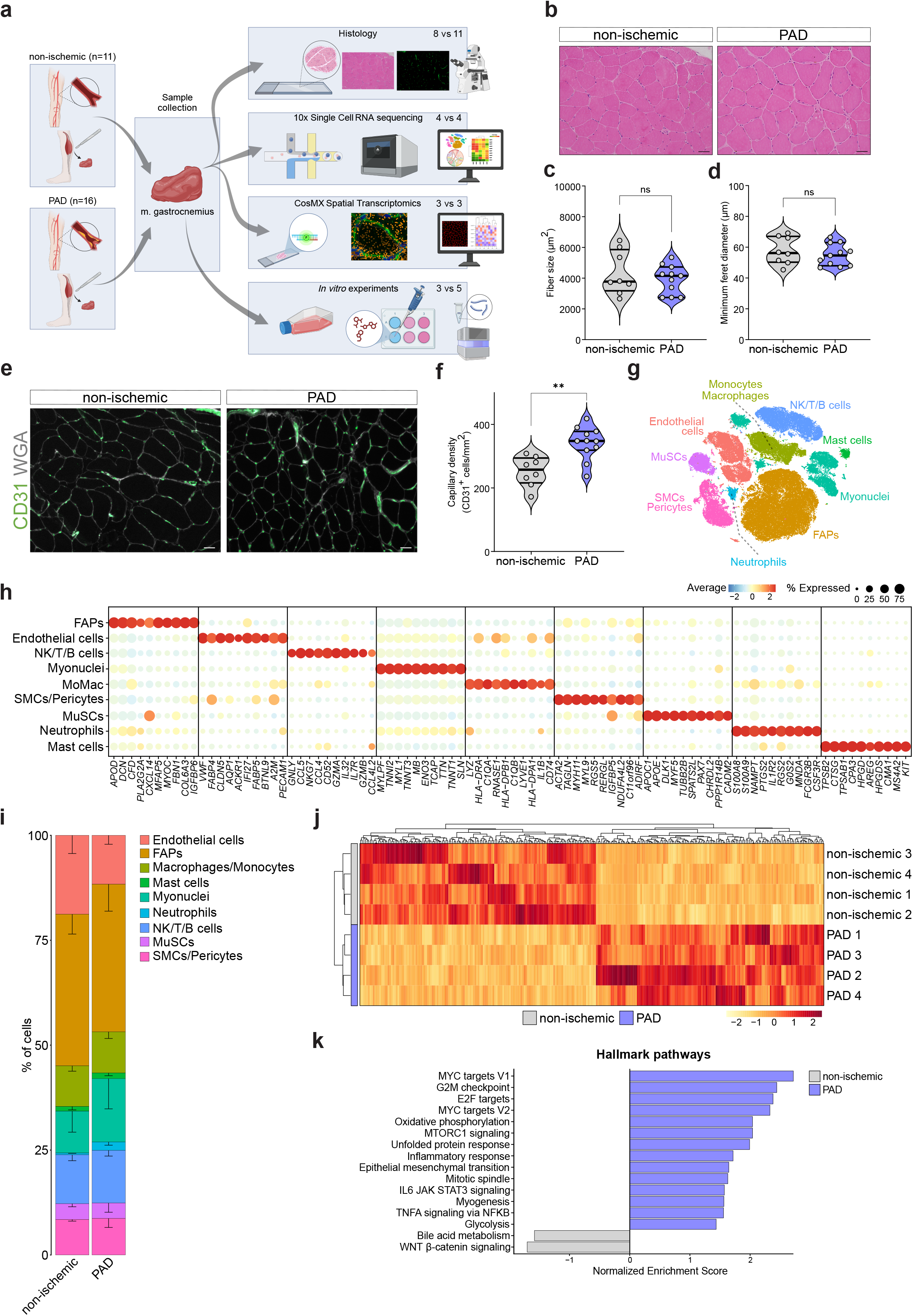
scRNAseq reveals cell heterogeneity in PAD. a, Graphical abstract of the experimental approach used in this study. **b**, Representative images of haematoxylin and eosin (H&E)-stained sections of gastrocnemius muscle from non-ischemic (n=8) and PAD (n=11) patients (Scale bar: 50 μm). **c,d**, Quantification of fiber size in μm^2^ (**c**) and minimum feret diameter in μm (**d**) from panel b. **e**, Representative images of gastrocnemius muscle from non-ischemic (n=8) and PAD (n=11) stained for the EC marker (CD31, green) and WGA (white) (Scale bar: 50 μm). **f**, Quantification of capillary density from panel e measured as the number of CD31^+^ cells per mm^2^. **g**, TSNE plot of cell populations identified from human gastrocnemius muscle, color-coded by the identified populations. **h**, Dot plot of centered logcounts values from cell type-specific marker genes. Color and size of the dots indicate the centered logcount value and the proportion of cells that express the gene, respectively. **i**, Stacked bar plots showing cluster percentage from non-ischemic (n=4) and PAD (n=4) patients, color-coded by cluster. Each stack represents mean - SEM. **j**, Heatmap of centered normalized values from top high variable genes (adjusted p-value < 0.05) in each patient based on pseudobulk analysis between non-ischemic and PAD conditions, color indicates the centered normalized values. **k**, Bar plots showing Normalized Enrichment Score of significant (adjusted p-value < 0.05) Hallmark pathways (MsigDB) from GSEA analysis on the pseudobulk dataset, color indicates the condition. Each dot represents a single patient in panels c,d and f. Student’s t test (two-tailed, unpaired, parametric, ns > 0.05, ^**^p < 0.01) was used in c,d and f. Wald test (as implemented in DESeq2 package) was used in j. Adaptive multi-level Monte-Carlo scheme (as implemented in fgsea package) was used in k. P = 0.4535 (**c**), P = 0.4585 (**d**), P = 0.0012 (**f**). Panel **a** created with BioRender.com.

**Table 1.**
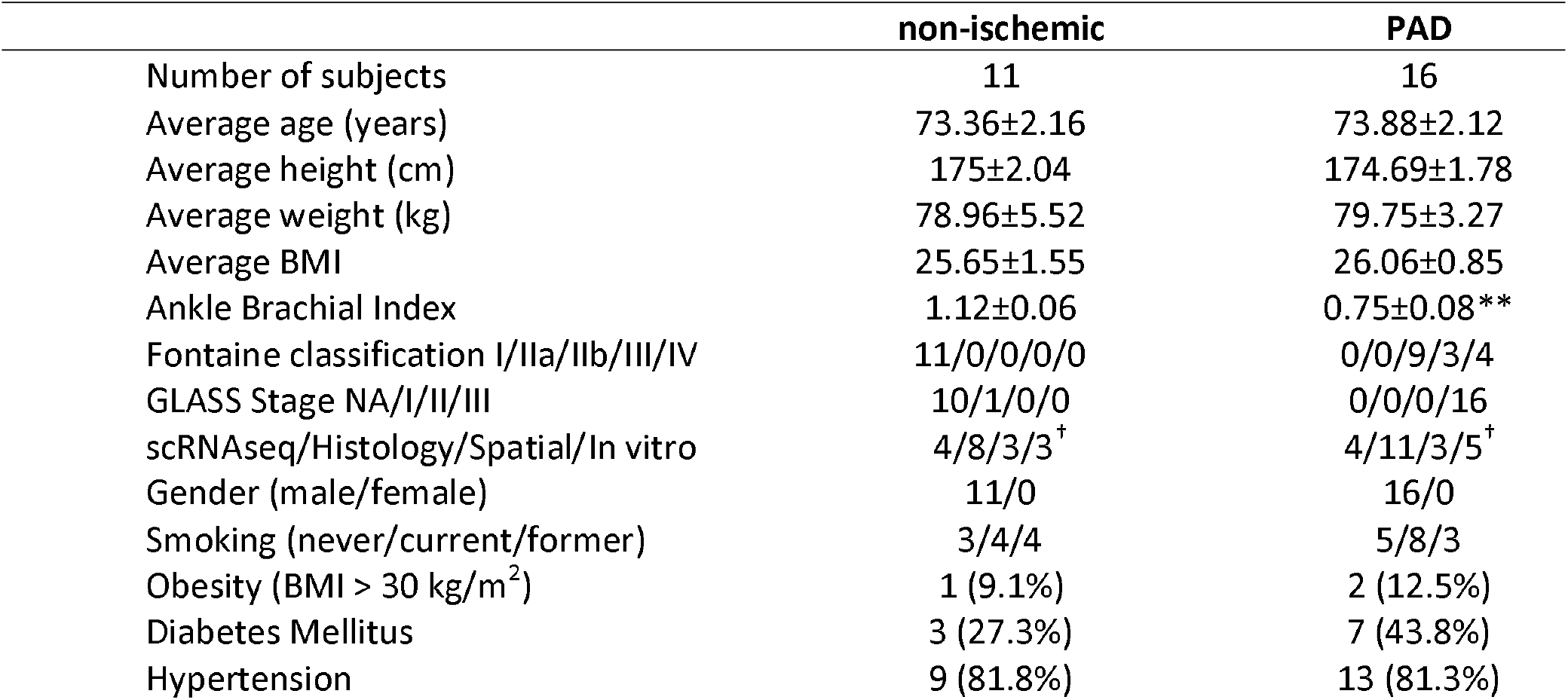
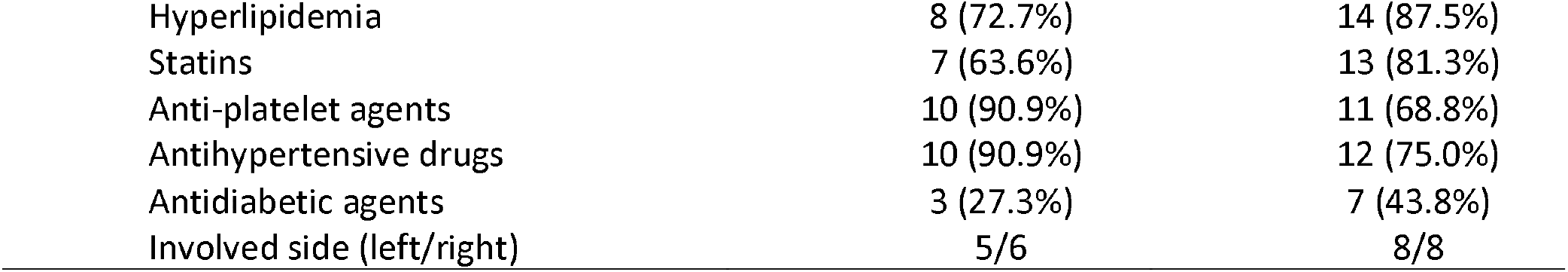
Patient demographics. Continuous values are presented as mean ± standard error of the mean. BMI indicates Body Mass Index. ^**^p < 0.01, Student’s t test (unpaired, two-tailed). ^†^A subset of histology samples was used for scRNAseq and/or spatial experiments; In vitro experiments were performed with 3 PAD samples for bulk RNAseq and 3 vs 5 for conditioned media experiments.

|  | non-ischemic | PAD |
| --- | --- | --- |
| Number of subjects | 11 | 16 |
| Average age (years) | 73.36±2.16 | 73.88±2.12 |
| Average height (cm) | 175±2.04 | 174.69±1.78 |
| Average weight (kg) | 78.96±5.52 | 79.75±3.27 |
| Average BMI | 25.65±1.55 | 26.06±0.85 |
| Ankle Brachial Index | 1.12±0.06 | 0.75±0.08** |
| Fontaine classification I/IIa/IIb/III/IV | 11/0/0/0/0 | 0/0/9/3/4 |
| GLASS Stage NA/I/II/III | 10/1/0/0 | 0/0/0/16 |
| scRNAseq/Histology/Spatial/In vitro | 4/8/3/3<sup>†</sup> | 4/11/3/5<sup>†</sup> |
| Gender (male/female) | 11/0 | 16/0 |
| Smoking (never/current/former) | 3/4/4 | 5/8/3 |
| Obesity (BMI > 30 kg/m<sup>2</sup>) | 1 (9.1%) | 2 (12.5%) |
| Diabetes Mellitus | 3 (27.3%) | 7 (43.8%) |
| Hypertension | 9 (81.8%) | 13 (81.3%) |
| Hyperlipidemia | 8 (72.7%) | 14 (87.5%) |
| Statins | 7 (63.6%) | 13 (81.3%) |
| Anti-platelet agents | 10 (90.9%) | 11 (68.8%) |
| Antihypertensive drugs | 10 (90.9%) | 12 (75.0%) |
| Antidiabetic agents | 3 (27.3%) | 7 (43.8%) |
| Involved side (left/right) | 5/6 | 8/8 |

Myofiber size nor diameter was affected in PAD patients (Fig. 1b-d). Also, we did not observe muscle necrosis but noticed a small non-significant increase in the percentage of regenerating fibers (Extended Data Fig. 1a) and increased CD31^+^ capillary (Fig. 1e-f) and arteriole densities (Extended Data Fig. 1b). We next performed scRNAseq in a subset of patients (Table 1) and generated a dataset comprising 106,566 cells (2359 genes/cell). Clustering revealed 9 major cell populations (Fig. 1g) that we manually annotated based on the expression of well-known marker genes^23^ (Fig. 1h, Fibro-adipogenic progenitors (FAPs), Natural killer (NK), Monocytes/Macrophages (MoMac), Smooth muscle cells (SMCs), Muscle stem cells (MuSCs)). We identified the main cell types in skeletal muscle (Fig. 1g-h) and did not detect obvious differences in cell proportions between groups (Fig. 1i and Extended Data Fig. 1c).

We additionally performed single-cell spatial transcriptomics to directly localize scRNAseq populations (Table 1). We identified muscle fibers and mononuclear cells by combining transcript detection with cell segmentation (Extended Data Fig. 1d) and annotated each cell through integration with the scRNAseq dataset (Extended Data Fig. 1e). Approximately 75% of the detected cells were fiber-derived (Extended Data Fig. 1f), reflecting the predominance of myonuclei in skeletal muscle. Despite these expected differences with the scRNAseq data, ECs, FAPs and SMCs/Pericytes still constituted most of the mononuclear fraction.

We first compared the scRNAseq patients by pseudobulk analysis, which showed different transcriptomic signatures between groups (Fig. 1j and Supplementary Data 1). Gene Set Enrichment Analysis (GSEA) revealed increased TNFA and IL6 signaling, as well as inflammatory activation, unfolded protein response and metabolic pathways, while WNT signaling and bile acid metabolism were repressed (Fig. 1k). Additionally, GSEA using a geneset collection from Human Phenotype Ontology (*Abnormality of muscle physiology, HP:0011804*) revealed increased PAD-like clinical symptoms while no term was enriched in non-ischemic patients (Extended Data Fig. 1g).

Next, we performed a comprehensive analysis on ECs, macrophages and SMCs/Pericytes because of their crucial role in angiogenesis, inflammation, and tissue repair^21^. The data including all identified cell populations can be interactively explored at https://shiny.debocklab.hest.ethz.ch/Turiel-et-al/.

### Human skeletal muscle contains *ATF3/4* endothelial subpopulations with immunoregulatory signatures

We selected ECs from the complete dataset and detected 6 different populations, which we annotated using described marker genes^24,25^ (Fig. 2a-b). Each population presented specific signatures, demonstrating clustering robustness (Fig. 2c and Supplementary Data 1). We identified two distinct populations of venous and capillary ECs that expressed similar markers (Fig. 2b) but showed different transcriptomic signatures (Fig. 2a,c). Differential expression analysis between both venous clusters detected a set of genes specifically enriched in venous 1 and capillary 1 populations (Extended Data Fig. 2a). Over-representation analysis (ORA)^26^ revealed that these genes are associated with recruitment and functional modulation of immune cells by ECs including TNF, IL-17 and NF-kappa B pathways^17,27,28^ as well as a lipid and atherosclerosis signaling (Fig. 2d). A deeper analysis of these pathways revealed that they are involved in leukocyte recruitment and activation, cell adhesion and cytokine signaling (Extended Data Fig. 2b-c). Recent studies^29^ have identified immunomodulatory ECs (IMECs), and the genes that define IMECs^29^ were enriched in venous 1 ECs (Extended Data Fig. 2d). We also observed higher expression in capillary 1 than in capillary 2 (Extended Data Fig. 2d, bottom panel), suggesting that both populations are ECs with immunoregulatory features.

**Fig. 2:**
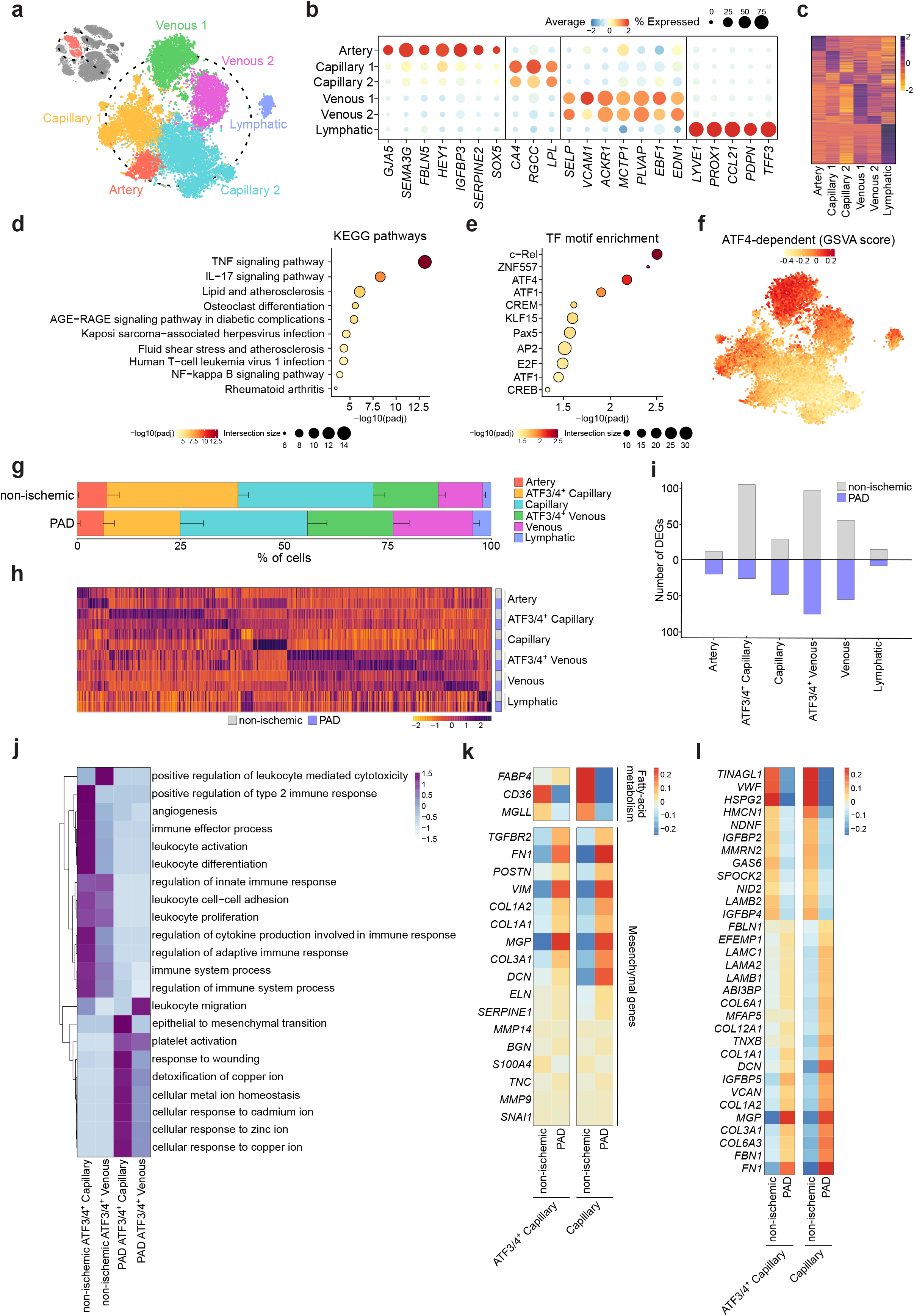
Human skeletal muscle contains several EC subtypes with different functional properties. **a**, TSNE plot of EC subtypes identified from human gastrocnemius muscle, color-coded by the identified subtypes. **b**, Dot plot of centered logcounts values from EC subtype-specific marker genes. Color and size of the dots indicate the centered logcount value and the proportion of cells that express the gene, respectively. **c**, Heatmap of centered logcounts of DEGs (adjusted p-value <0.05) between EC subtypes, color indicates the centered logcount value. **d**,**e**, ORA analysis from genes upregulated in Venous 1 compared to Venous 2 (see Extended Data Fig. 2a) using KEGG pathways (**d**) or TF motif enrichment (**e**), color and size of the dots indicates the -log10 adjusted p-value and the number of genes in each category (intersection size), respectively. **f**, TSNE plot of GSVA scores using the ATF4 dependent geneset from Fan et al (see Methods), color-coded by the GSVA score. **g**, Stacked bar plots showing cluster percentage from non-ischemic (n=4) and PAD (n=4) patients, color-coded by cluster. Each stack represents mean - SEM. **h**, Heatmap of centered logcounts of DEGs (adjusted p-value < 0.05) specific for each EC subtype in each condition, color indicates the centered logcount value. **i**, Bar plots showing the number of DEGs for each EC subtype in each condition, color indicates the condition. **j**, Heatmap of centered values from ORA analysis over the DEGs in *ATF3/4*^+^ venous and capillary ECs between conditions, color indicates the centered values. **k,l**, Heatmap of centered logcounts values of fatty acid metabolism and mesenchymal markers as defined in Tombor et al (**k**) and core matrisome genes as defined in Hynes et al. (**l**), color indicates the centered values. Wilcoxon Rank Sum test (as implemented in Seurat package) was used in c and h. Fisher’s one-tailed test (as implemented in g:Profiler) was used in d and e.

ORA for transcription factor (TF) motif enrichment identified c-Rel, ZNF557 and ATF4 as top TFs regulating the genes enriched in both venous 1 and capillary 1 populations (Fig. 2e and Extended Data Fig. 2a). We also applied SCENIC^30^, which generates a *TF regulon* activity score based on the coexpression of TFs and their target genes. SCENIC did not detect ZNF557 as an active TF, due to its few detected target genes (intersection size in Fig. 2e), and c-Rel activity was relatively weak (Extended Data Fig. 2e). In contrast, ATF4-dependent TFs factors showed high and specific activity in venous 1 and capillary 1 clusters (Extended Data Fig. 2e), so we focused on *ATF3/4*. We previously identified *Atf3/4^+^* muscle ECs in mouse that control exercise-induced angiogenesis^24^. The marker genes defining *Atf3/4^+^* ECs were highly enriched in both venous 1 and capillary 1 human clusters (Extended Data Fig. 2f). Conversely, the marker genes from the human populations (Extended Data Fig. 2a) were also controlled by ATF4 in ECs (Extended Data Fig. 2g). We also used an ATF4-dependent gene signature to perform gene set variation analysis (GSVA)^31^ which calculates a score for each cell based on the expression levels of an input gene signature. The ATF4-dependent GSVA score was enriched in venous 1 and capillary 1 clusters (Fig. 2f) and we therefore annotated these populations as *ATF3/4^+^* venous ECs and *ATF3/4^+^* capillary ECs. The fraction of *ATF3/4^+^* capillaries was reduced in PAD when compared to non-ischemic samples (Fig. 2g). We also identified all EC subtypes in the spatial dataset and found very consistent cluster proportions between both technologies (Extended Data Fig. 2h-i).

We next analyzed the functional contribution of *ATF3/4^+^* ECs in PAD by using a preclinical mouse model. We assessed muscle reperfusion following hindlimb ischemia (HLI) in wild-type (WT) and EC-specific inducible ATF4 knockout mice (*Atf4*^ECKO^). HLI efficiently reduced blood flow in both genotypes leading to similar levels of ischemia-induced damage 3 days after surgery (Extended Data Fig. 3a,b), characterized by large degenerative areas with cell infiltration and regenerating fibers. *Atf4*^ECKO^ mice exhibited impaired blood flow recovery starting 7 days after HLI (Extended Data Fig. 3c,d), which coincided with reduced vessel diameter and density (Extended Data Fig. 3e-g) and lower EC proliferation (Extended Data Fig. 3h). As a result, *Atf4*^ECKO^ mice showed larger damaged muscles 7 and 14 days after surgery (Extended Data Fig. 3a,b), illustrating the functional importance of ATF3/4^+^ ECs to ischemia-induced revascularization in a preclinical model of PAD.

### ECs undergo profound transcriptomic rewiring during PAD

Next, we assessed the transcriptional differences between conditions by calculating the number of differentially expressed genes (DEGs) that are specific for each EC population (Fig. 2h and Supplementary Data 1). *ATF3/4^+^* venous and *ATF3/4^+^* capillary ECs showed the highest number of DEGs between non-ischemic and PAD samples (Fig. 2i). ORA on the DEGs in *ATF3/4^+^* venous or *ATF3/4^+^* capillary ECs (Extended Data Fig. 4a and Supplementary Data 1) revealed that few enriched processes were activated in PAD (Extended Data Fig. 4b and Supplementary Data 1), while most of them were downregulated indicating that many functions exerted by these populations are repressed, including angiogenesis, immune regulation, leukocyte interactions as well as metabolism (Fig. 2j and Extended Data Fig. 4c). Conversely, few gene sets related to epithelial-to-mesenchymal transition (EMT), platelet activation, wound healing, and cellular responses to metal ions were activated in PAD (Fig. 2j).

The activation of EMT in *ATF3/4^+^* capillary ECs (Fig. 2j) prompted us to EndoMT. Focusing on capillary ECs, we noticed EndoMT features^32^, including repressed fatty acid metabolism genes and partial mesenchymal activation (Fig. 2k). Some ‘canonical’ EndoMT genes like *SNAI1, S100A4* and *SERPINE1* were not affected in PAD while several extracellular matrix (ECM) components, such as *COL1A1, COL1A2* and *FN1* were highly upregulated (Fig. 2k). This phenotype was present in both capillary subpopulations and they also lowered *CDH5* expression but did not completely lose it (Extended Data Fig. 4d), suggesting partial EndoMT. We then investigated how ECs rewire their matrisome, a set of genes that define ECM composition and function^33^. Capillary ECs in PAD only changed 11.7% (32/274) of the core matrisome genes but showed significant activation of ECM components such as collagens, *FN1, FBN1*, as well as alterations in components of the basement membrane such as laminins, *HSPG2*, and *NID2* (Fig. 2l). Immunohistological analysis confirmed increased expression of the mesenchymal marker fibronectin (FN1) by CD31^+^ ECs in PAD (Extended Data Fig. 4e,f). Taken together, ECs in PAD undergo profound transcriptomic alterations that impair angiogenesis and immunoregulatory functions but promote EndoMT.

Finally, we used DEGs between PAD and non-ischemic ECs to predict drugs^34^ that can reverse the PAD EC phenotype. This analysis predicted a set of compounds with this therapeutic potential (Fig. 3a), including withaferin-a or saracatinib, drugs that target inflammation and endothelial dysfunction^35,36^. We focused on celastrol for its ability to improve vascular remodeling and to reduce endothelial dysfunction^37^. We treated primary isolated muscle ECs (Extended Data Fig. 4g, see Methods) from 3 PAD patients for 24h with vehicle (DMSO) or celastrol (125 or 250 nM^38^). RNAseq analysis showed a dose-dependent effect across patients (Fig. 3b-c and Supplementary Data 1), including upregulated (C1 in Fig. 3c) and downregulated genes (C2 and C3). ORA (Fig. 3d) confirmed celastrol-dependent upregulation of processes related to cellular homeostasis, detoxification and response to metal ions (C1) while downregulating EMT, response to wounding (C2), leukocyte migration and inflammation (C2 and C3), indicating the ability of celastrol to partially reverse the PAD phenotype observed in the scRNAseq dataset (Fig. 2j). Celastrol also activated ATF4 or glucosamine signatures (Fig. 3e), which improves perfusion recovery in PAD preclinical models by activating ATF4^39^. These findings suggest the potential of our dataset for identifying compounds that could potentially revert PAD phenotypes.

**Fig. 3:**
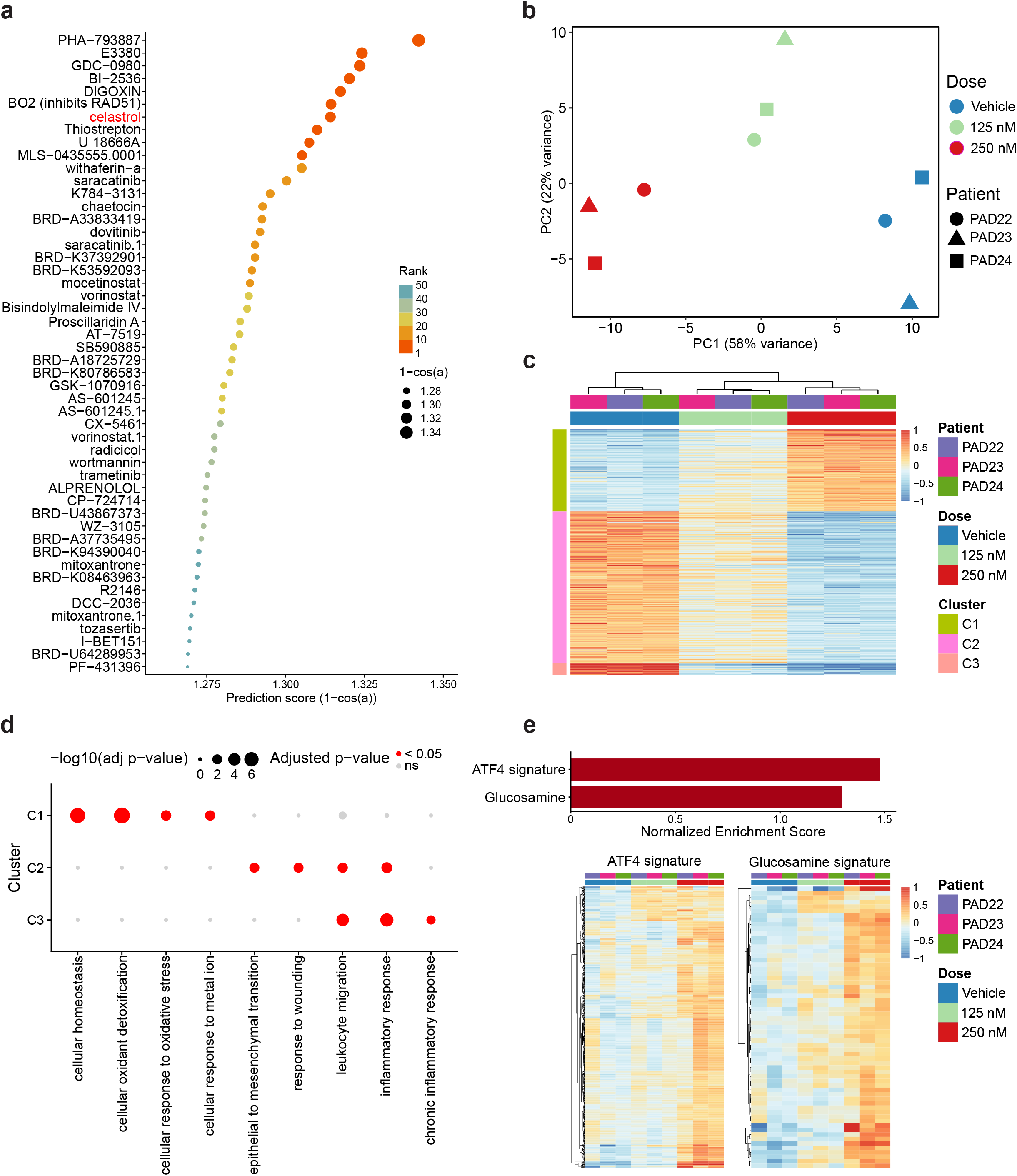
Celastrol partially reverses the EC PAD signature. **a,** Dot plot of prediction scores (derived from L1000CDS^2^ search engine) from different compounds with the potential to reverse EC PAD signature. Color and size of the dots indicate the rank and the prediction score, respectively. **b,** Principal Component Analysis (PCA) showing sample distances between different ECs (isolated from PAD patients) treated with Vehicle (DMSO) or celastrol (125 or 250 nM). Color and shape indicate treatment and patient of origin, respectively. **c,** Heatmap showing centered normalized values of DEGs (adjusted p-value < 0.05) between Vehicle and celastrol 250 nM, color indicates the centered normalized value. Genes (rows) are organized and color-coded based on clustering of gene expression profiles. **d,** ORA analysis of the gene clusters from panel c, color and size indicates adjusted p-values. **e,** Bar plot (top panel) showing Normalized Enrichment Score (adjusted p-value < 0.05) from GSEA using ATF4 (Fan et al) and glucosamine signatures (Alhusban et al). Heatmap (bottom panel) showing centered normalized values of top enriched genes from each signature, color indicates the centered normalized value. Wald test (as implemented in DESeq2 package) was used in c. Fisher’s one-tailed test (as implemented in g:Profiler) was used in d. Adaptive multi-level Monte-Carlo scheme (as implemented in fgsea package) was used in e.

### *LYVE1*^hi^ MHCII^low^ macrophages are the dominant macrophage population in PAD

In experimental PAD models, monocyte-derived macrophages contribute to vascularization^16,19,21,40–42^. However, acute mouse models do not reflect chronic scenarios, so the contribution of macrophages to human PAD is poorly understood. We decided to further investigate the Monocytes/Macrophages cluster (MoMac), selecting these cells from the complete dataset where we identified and annotated 5 distinct populations^43,44^ (Fig. 4a-c and Supplementary Data 1). This dataset included classical monocytes (cMonocytes), *CD16*(*FCGR3A*)^+^ monocytes, dendritic cells (DCs) and two macrophage populations. Alternatively, we could also classify the populations as *CCR2*^+^C*X3CR1*^low^*CD14*^hi^*CD16*^low^ (cMonocytes and DCs) and *CCR2*^-^*CX3CR1*^hi^*CD14*^low^*CD16*^hi^ (*CD16*^+^ Monocytes)(Extended Data Fig. 5a).

**Fig. 4:**
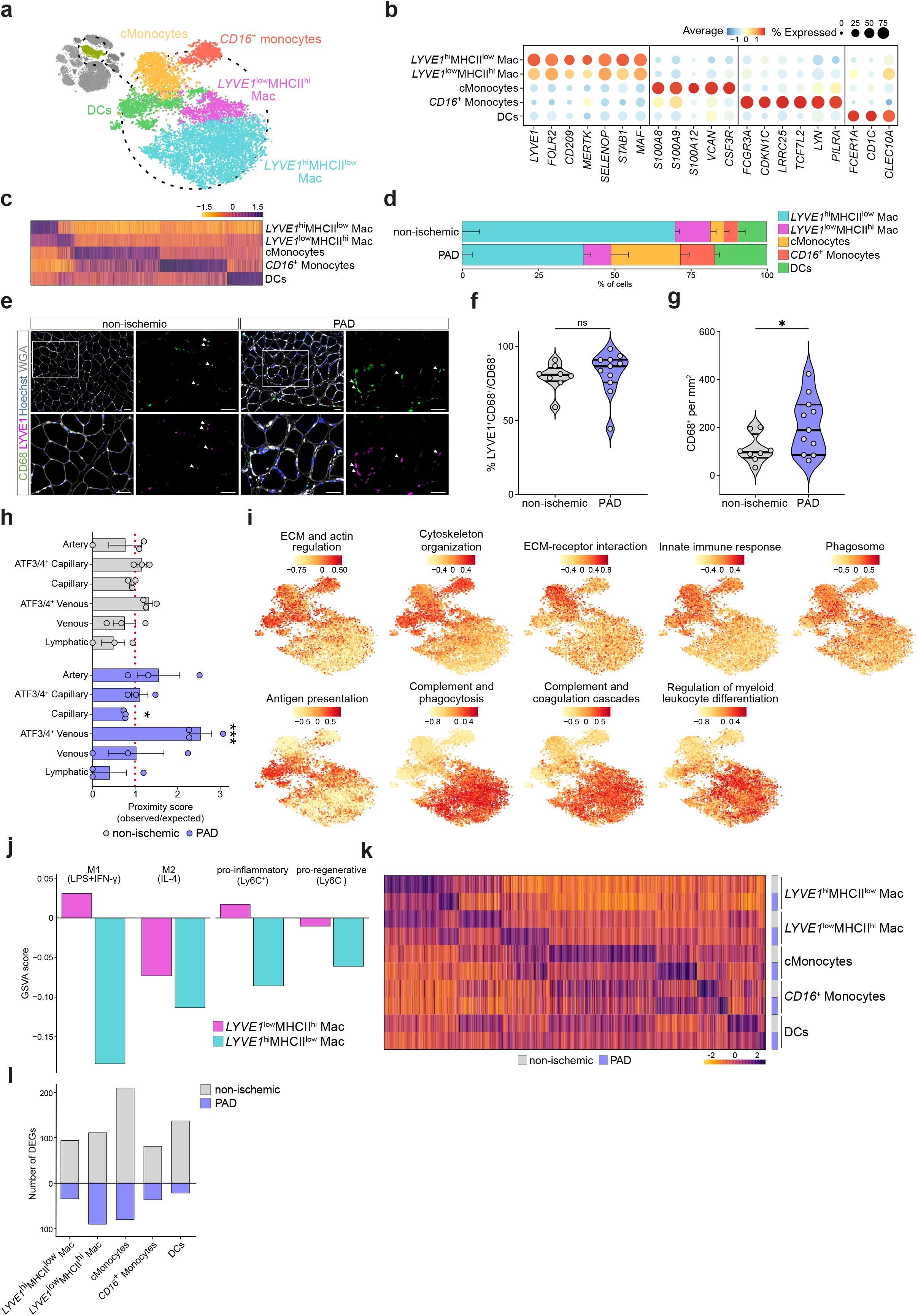
*LYVE1*^hi^MHCII^low^ macrophages are the dominant macrophage population in PAD. **a**, TSNE plot of Monocytes/Macrophages subtypes identified from human gastrocnemius muscle, color-coded by the identified subtypes. **b**, Dot plot of centered logcounts values from MoMac subtype-specific marker genes. Dot color indicates centered logcounts, and size reflects the proportion of cells expressing the gene. **c**, Heatmap of centered logcounts of DEGs (adjusted p-value <0.05) between MoMac subtypes, color indicates the centered logcount value. **d**, Stacked bar plots showing cluster percentage from non-ischemic (n=4) and PAD (n=4) patients, color-coded by cluster. Each stack shows mean - SEM. **e**, Representative images of gastrocnemius muscle from non-ischemic (n=8) and PAD (n=11) stained for a pan-macrophage marker (CD68, green), LYVE1 (magenta), cell nuclei (Hoechst, blue) and WGA (white). Top left panel has a lower magnification compared to the other 3 pictures. White rectangle indicates the zoomed area displayed in the remaining panels (Scale bar: 50 μm). **f-g**, Quantification of the percentage of LYVE1^+^CD68^+^ cells from the total of CD68^+^ cells (**f**) and the total number of macrophages measured as the number of CD68^+^ cells/mm^2^ (**g**). **h**, Bar plot showing proximity scores (see Methods) of *LYVE1*^hi^MHCII^low^ macrophages to each EC subtype in the spatial dataset from non-ischemic (n=3) and PAD (n=3) patients. Color coded by condition. Each bar represents mean ± SEM. **i,** TSNE plots showing GSVA scores for different macrophage activation processes as defined by Sanin et al, color-coded by GSVA score. **j**, GSVA scores from different pro-inflammatory and pro-regenerative signatures as defined by Varga et al between *LYVE1*^hi^ and *LYVE*^low^ macrophages, color-coded by cluster. **k**, Heatmap of centered logcounts of DEGs (adjusted p-value < 0.05) specific for each MoMac subtype in each condition, color indicates the centered logcount value. **l**, Bar plot showing the number of DEGs for each MoMac subtype in each condition, color indicates the condition. Each dot represents a single patient in panels f, g and h. Wilcoxon Rank Sum test (as implemented in Seurat package) was used in c and k. Student’s t test (two-tailed, unpaired, parametric, ns > 0.05) was used in f and Mann-Whitney U test (two-tailed, unpaired, non-parametric, ^*^p < 0.05). was used in g. Binomial test (one-tailed, observed vs expected, ^*^p < 0.05) was used in h. P = 0.3511 (**f**), P = 0.0473 (**g**), P = 0.0066 (**h**, PAD Capillary), P = 0.0007 (**h**, PAD *ATF3/4*^+^ Venous).

Macrophage subpopulations were annotated as *LYVE1*^hi^MHCII^low^*CX3CR1*^low^ and *LYVE1*^low^MHCII^hi^*CX3CR1*^hi^ (Extended Data Fig. 5b), matching previous characterization^45^. In mouse, *LYVE1*^hi^MHCII^low^ macrophages often have a perivascular location and a distinct ontology, phenotype and function compared to monocyte-derived macrophages^45^. *LYVE1*^hi^MHCII^low^ macrophages in our dataset expressed high levels of tissue-resident macrophage markers including *LYVE1, FOLR2, CD209, MERTK, SELENOP, STAB1* and *MAF*^44,46,47^ (Fig. 4b). *LYVE1*^hi^ macrophages were the largest macrophage population, even though their percentage was lower in PAD (70±6% and 40±3%, respectively, Fig. 4d). LYVE1/CD68 co-staining confirmed the predominance of *LYVE1*^hi^ macrophages in both conditions (Fig. 4e,f), despite a significant increase in CD68^+^ monocyte/macrophages in PAD muscle (Fig. 4g), consistent with previous work^21^. The spatial dataset also showed *LYVE1*^hi^ macrophages as the main MoMac population in human muscle (Supplementary Fig. 5c). Notably, the percentage of *LYVE1*^+^ macrophages over the total fraction of MoMac cells (CD68^+^) was similar in both spatial and immunofluorescence approaches (≈80-85%, Extended Data Fig. 5c and Fig. 4f) and was not affected in PAD in none of them. Since the scRNAseq dataset could include (due to residual blood in the sample) circulating MoMac populations such as cMonocytes and DCs, the proportional reduction in *LYVE1*^+^ macrophages was likely secondary to increased presence of those populations in PAD (Fig. 4d). Moreover, the increase of cMonocytes (Fig. 4d), but not monocyte-derived *LYVE1*^low^MHCII^hi^ macrophages, suggests impaired monocyte-to-macrophage differentiation in PAD, which was confirmed using ORA (Extended Data Fig. 5d), or could be secondary to increased cMonocyte numbers (Fig. 4d).

To evaluate whether perivascular *LYVE1*^hi^ macrophages^45^ preferentially localize to specific vessels, we calculated a proximity score using spatial transcriptomics, defined as the ratio of observed versus expected *LYVE1*^hi^MHCII^low^ macrophages closest to a given EC subtype. A score >1 indicates that *LYVE1*^hi^MHCII^low^ macrophages are usually closer to a specific EC subtype than expected by chance. In non-ischemic samples, we did not observe any spatial enrichment of *LYVE1*^hi^MHCII^low^ macrophages (Fig 4h). However, in PAD, they exhibited a significant proximity score to ATF3/4^+^ Venous (Fig. 4h), indicating a preferential localization to these ECs.

MoMacs exert many functions depending on the biological context. Sanin et al^48^ categorized macrophage activation into four functional pathways: phagocytic, oxidative stress, inflammatory and remodeling (regenerative). We utilized this framework to characterize the functional differences between the MoMac populations in our dataset. *CD16*^+^ Monocytes, cMonocytes and DCs were characterized by processes related to early stage activation (ECM/cytoskeleton regulation) and inflammatory/oxidative paths (innate immune response and phagosome) (Fig. 4i). *LYVE1*^low^ macrophages and DCs, showed a profile of antigen presentation, consistent with their higher MHCII levels (Extended Data Fig. 5b) and with their previous characterization^45^. Both *LYVE1*^hi^ and *LYVE1*^low^ macrophages were characterized by a specific phagocytic profile as they were enriched in processes related to complement signaling, phagocytosis and regulation of leukocyte differentiation.

The differential and overlapping functions of *LYVE1*^hi^ versus *LYVE1*^low^ macrophages upon loss of muscle homeostasis, including PAD, are poorly understood, so we explored whether they present pro-inflammatory (M1) or anti-inflammatory, pro-regenerative (M2)^49^ macrophage signatures. We used published genesets associated with M1 or M2 macrophages^50^ under *in vitro* stimulation as well as *in vivo* states of macrophages isolated from cardiotoxin-injured muscles (Ly6C^+^ macrophages at day 2 after injury (pro-inflammatory) and Ly6C^-^ at day 4 after injury (pro-regenerative)). *LYVE1*^hi^ macrophages expressed lower levels of both M1/inflammatory and M2/regenerative signatures (Fig. 4j), consistent with their proposed ‘homeostatic’ function. Additionally, *LYVE1*^low^ macrophages showed higher expression of both signatures, although they exhibited a more pronounced pro-inflammatory signature (Fig. 4j). Thus, *LYVE1*^hi^ macrophages do not reflect the genetic fingerprint of M1/inflammatory nor M2/regenerative macrophages.

### *LYVE1*^hi^ MHCII^low^ macrophages get activated during PAD

To assess whether MoMac populations are altered upon PAD, we detected DEGs for every individual population and observed alterations across all MoMac populations (Fig. 4k,l and Supplementary Data 1). Although cMonocytes showed the highest number of DEGs, most of their biological processes were downregulated in PAD, including terms related to leukocyte/macrophage differentiation (Extended Data Fig. 5d). Since *LYVE1*^hi^MHCII^low^ macrophages are the dominant population, and their role in muscle is poorly understood, we focused on those. Most DEGs in *LYVE1*^hi^MHCII^low^ macrophages were upregulated during PAD (Extended Data Fig. 5e and Supplementary Data 1) and associated to inflammatory and immune responses, cytokine production and macrophage activation (Extended Data Fig. 5f-g and Supplementary Data 1). The top repressed processes in PAD were related to heat responses and protein folding (Extended Data Fig. 5g) including genes such as *HSPA1A* and *HSPA1B*, which have been associated with atheroprotective and anti-inflammatory roles in macrophages^51,52^. Altogether, *LYVE1*^hi^MHCII^low^ macrophages get activated and acquire pro-inflammatory properties during PAD.

### Disruption of cellular communication networks in the muscle microenvironment during PAD

To study cell-cell communication, we applied CellChat^53^ to the complete scRNAseq dataset. During PAD, the number of interactions slightly increased (Fig. 5a), although the strength of these interactions was generally reduced (Fig. 5b). The increased number of interactions was mainly mediated by ECs, SMcs/Pericytes, FAPs and MuSCs (Fig. 5c). Only neutrophils showed an increase in interaction strength with MoMac populations during PAD (Fig. 5d). FAPs were the main signal senders in the non-ischemic muscle microenvironment whereas MoMac populations were the main receptors (Fig. 5e). In PAD, most populations reduced their interaction strength (Fig. 5e), suggesting disrupted communication in the PAD muscle microenvironment. Indeed, most pathways were reduced in PAD (Fig. 5f). MHC-II signaling was one of the top reduced communications (Fig. 5f) with MoMac and ECs as they main cell types involved (Extended Data Fig. 6a). ECs reduced VEGF signaling in PAD (Extended Data Fig. 6b). MuSCs showed reduced NCAM, a cell adhesion molecule that marks MuSCs, interactions with FAPs (Extended Data Fig. 6c). ECs and *LYVE1*^+^ macrophages reduced PDGF signaling with FAPs (Extended Data Fig. 6d), a pathway involved in muscle hypertrophy and regeneration^54^. *ATF3/4*^+^ Venous showed reduced TRAIL signaling (Extended Data Fig. 6e), a key mediator in generating stable vessels in preclinical PAD models^55^. ECs also reduced interactions of several tight and adherens junctional genes including JAM, ESAM and CDH5 (Extended Data Fig. 6f-h). On the other hand, pro-inflammatory IL1 and IL6 were enriched in PAD, with the latter being specifically produced by *ATF3/4*^+^ Venous ECs (Extended Data Fig. 6i,j). All detected interactions and ligand-receptor pairs are available in Supplementary Data 1.

**Fig. 5:**
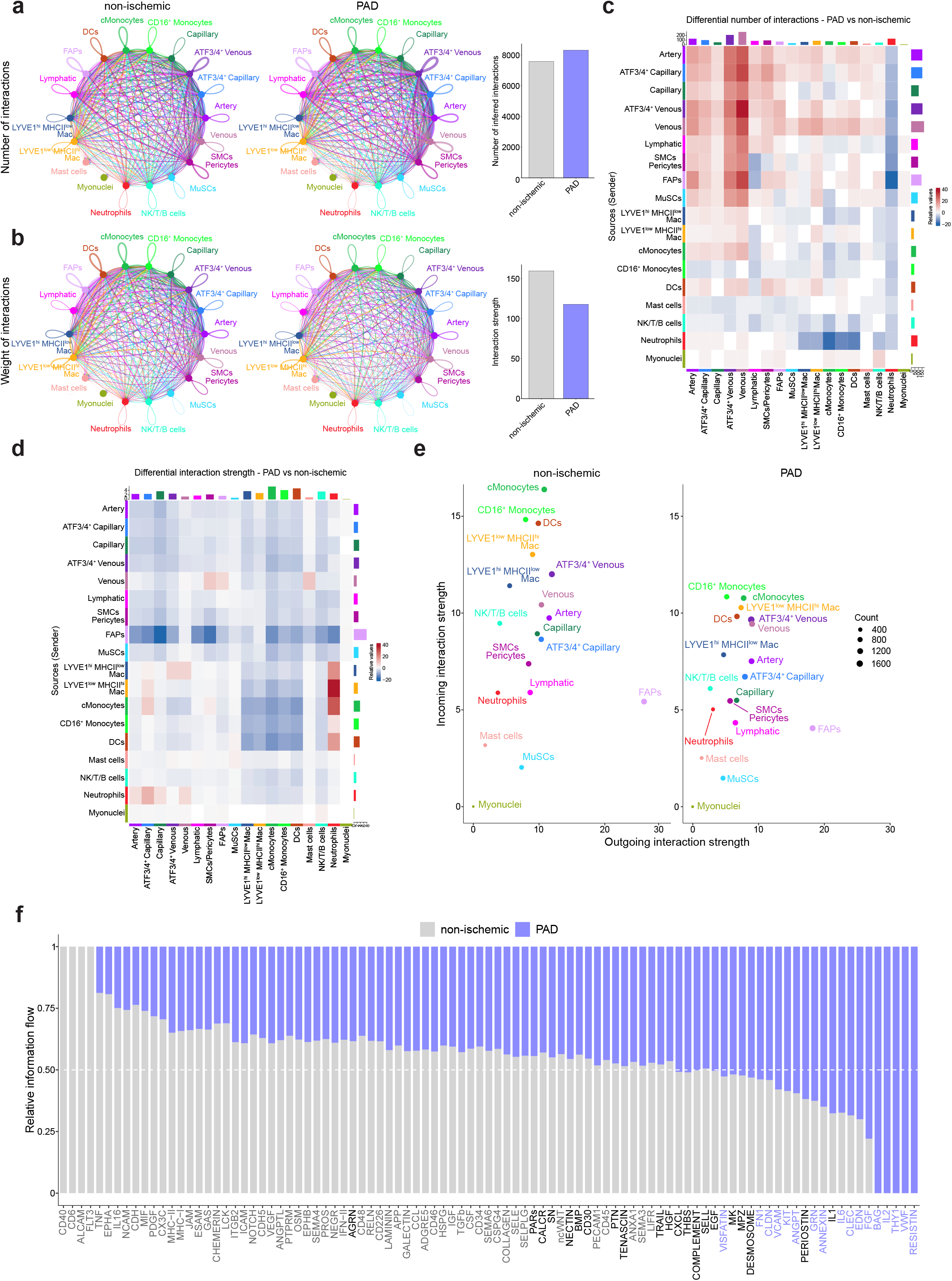
Cellular communication in human skeletal muscle is disrupted in PAD. **a-b**, Circle plots showing the number (**a**) and weight (**b**) of inferred interactions between cell types in non-ischemic and PAD (left and middle plots) and bar plots quantifying these parameters (right plots). Color and width of the edges indicates the sender and number or weight of interactions, respectively. **c-d**, Heatmaps showing increased (red) or decreased (blue) number (**c**) and weight (**d**) of inferred interactions in PAD. Labels on the left side indicate the cell type expressing the ligands and bottom labels the cell types receiving them. **e**, Scatter plots showing the strength of the outgoing (x-axis) and incoming (y-axis) interactions for each cell type in non-ischemic (left) and PAD (right). Color and size of the dots indicate the cell type and the number of inferred interactions, respectively. **f**, Bar plots showing the relative information flow for significant communication pathways between non-ischemic and PAD

### Altered SMC/Pericyte-Endothelium interactions suggest impaired vessel homeostasis during PAD

SMCs/Pericytes are essential for proper vessel function and stability. We therefore investigated how they interact with ECs during PAD. First, we annotated them based on well-known marker genes of both populations^56^ (Fig. 6a). Although they were present in similar proportions in non-ischemic samples (Fig. 6b), SMCs comprised around 75% of mural cells in PAD. Most DEGs were downregulated during PAD (Fig. 6c and Supplementary Data 1) but they increased pathways related to muscle development and contractility (Fig. 6d,e) while downregulating blood vessel and vasculature development processes.

**Fig. 6:**
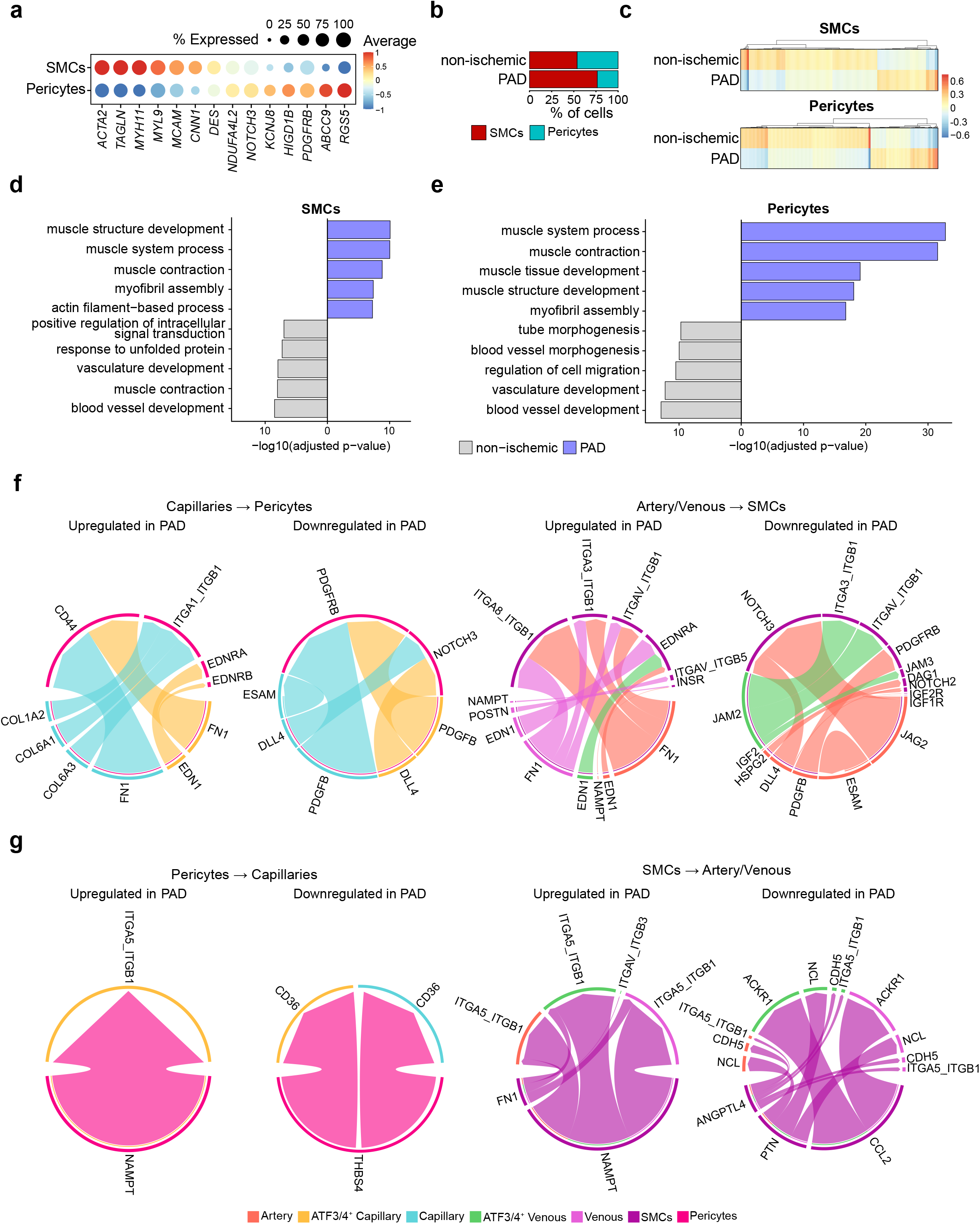
Cellular communication between mural cells and ECs indicates reduced vessel homeostasis. **a,** Dot plot of centered logcounts values from SMCs and Pericytes marker genes. Color and size of the dots indicate the centered logcounts values and the proportion of cells that express the gene, respectively. **b,** Stacked bar plots showing cluster percentage in each condition, color-coded by cluster. Each stack represents mean – SEM. **c,** Heatmap of centered logcounts of DEGs (adjusted p-value <0.05) from SMCs (top) and Pericytes (bottom) between conditions, color indicates the centered logcount value. **d-e,** ORA analysis of top biological processes of DEGs from panel c in SMCs (**d**) and Pericytes (**e**), color indicates in which condition the process is significant (adjusted p-value < 0.05). **f-g**, Circle plots of ligand and receptors up/downregulated in PAD from ECs to mural cells (**f**) and from mural cells to ECs (**g**). Color and width of the edges indicates the sender and weight of interactions, respectively. Wilcoxon Rank Sum test (as implemented in Seurat package) was used in c. Fisher’s one-tailed test (as implemented in g:Profiler) was used in d and e.

CellChat to study capillary-pericyte and artery/venous-SMC interactions revealed that ECs mainly increased FN1 signaling to both pericytes and SMCs populations (Fig. 6f), while capillaries and arteries reduced PDGFB-PDGFRB and JAG2/DLL4-NOTCH3 signaling, key mediators of recruitment and attachment to the endothelium^57,58^. Conversely, SMCs/Pericytes increased NAMPT signaling to ECs during PAD (Fig. 6g) but reduced THBS4 (pericytes) and *PTN, CCL2* and *ANGPTL4* (SMCs) communication (Fig. 6g). Together, these changes suggest a reduced capacity of SMC/Pericytes to maintain vessel homeostasis and endothelial attachment in PAD.

### Endothelial-macrophage crosstalk alters signaling networks driving inflammation and immune modulation in *LYVE*^hi^ MHCII^low^ macrophages during PAD

Finally, we analyzed EC-MoMac reciprocal communication, including Neutrophils due to their increased interaction strength with MoMacs during PAD (Fig. 5d). ECs upregulated FN1 interactions with all MoMacs while reducing MHCII signaling (Extended Data Fig. 7a). MoMacs increased NAMPT, linked to pro-inflammatory markers in CAD patients^59^, and *THBS1* (cMonocyte-specific) signaling, elevated in PAD patients and with anti-angiogenic effects in ECs^60^. Conversely, they downregulated *VEGFA* and *ANXA1* signaling (important for inflammatory resolution in muscle^61^) with ECs and Neutrophils, respectively (Extended Data Fig. 7b). Neutrophils upregulated *NAMPT* and *OSM*, a cytokine promoting EC inflammation and neutrophil recruitment^62,63^, while downregulating MHCII genes and *ANXA1*, further suggesting impaired inflammatory resolution (Extended Data Fig. 7c).

Next, we applied NicheNet^64^, which can additionally link ligand-receptor pairs to their downstream signaling networks, to further study EC-macrophage interactions in PAD. Since *ATF3/4*^+^ ECs present an immunomodulatory profile, and *ATF3/4*^+^ Venous were the closest to *LYVE1*^hi^MHCII^low^ macrophages in PAD (Fig. 4h), we focused on the communication between those populations. NicheNet identified different ligand-receptor pairs specifically downregulated (Fig. 7a) or upregulated (Fig. 7b) in PAD samples, indicating a profound rewiring of the communication network between *ATF3/4*^+^ ECs and *LYVE1*^hi^MHCII^low^ macrophages. Some of the top reduced signals during PAD were related to *GAS6* and *CSF1* signaling (Fig. 7a), while *IL6* signaling was enriched (Fig. 7b). We also used NicheNet to detect which upregulated genes in *LYVE1*^hi^MHCII^low^ macrophages during PAD could be downstream targets of ligands produced by *ATF3/4*^+^ ECs (Fig. 7c). These target genes included *CCL18, SGK1, PLAUR, CCL3, LMNA, HBEGF, IGFS6, FTH1* and *CTSB* (Fig. 7c) and are associated with migration, cytokine response, inflammatory response, and immune related processes (Fig. 7d).

**Fig. 7:**
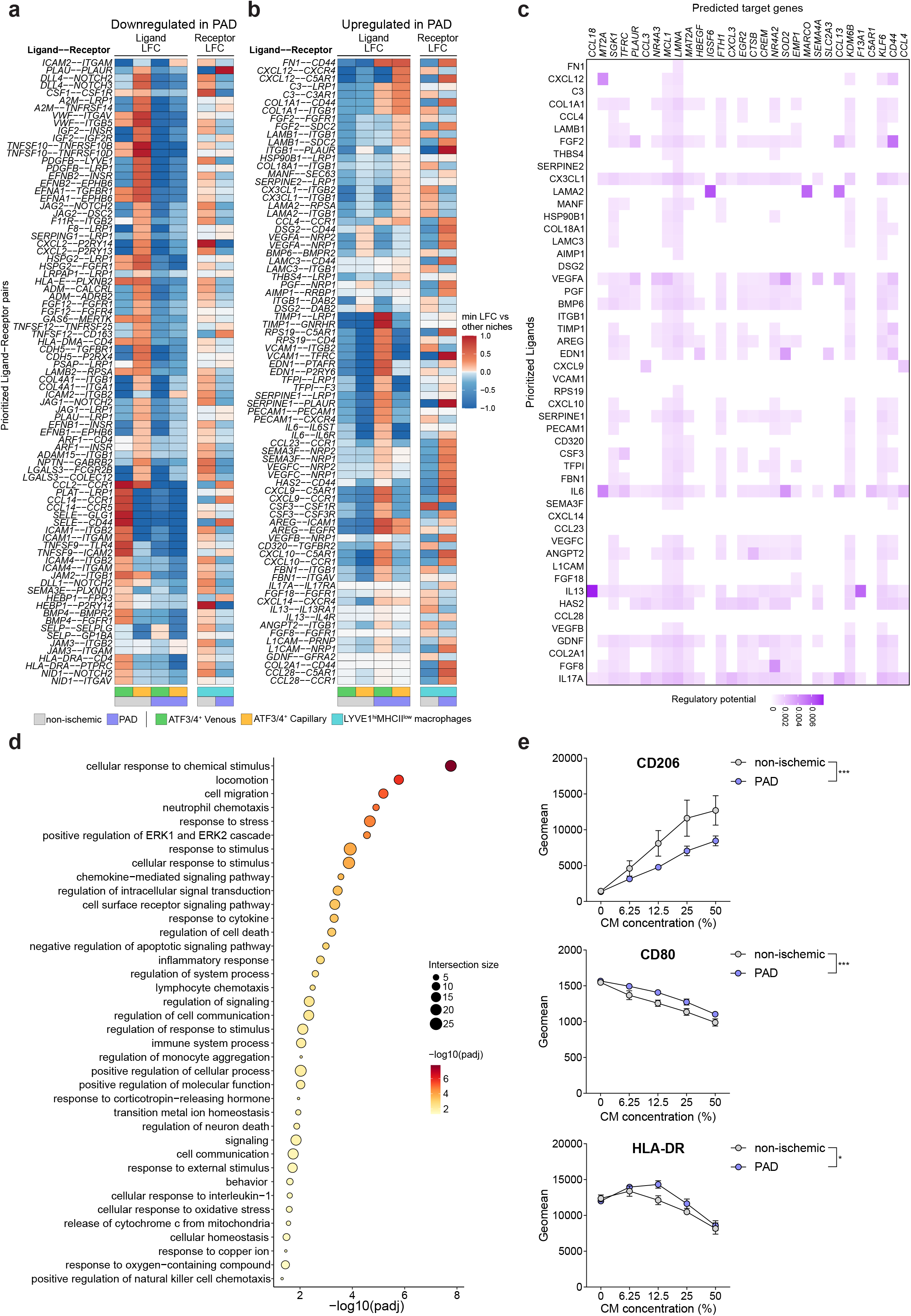
Cellular communication between *ATF3/4*^+^ ECs and *LYVE1*^hi^MHCII^low^ macrophages is altered during PAD. **a-b**, Heatmaps showing downregulated (**a**) and upregulated (**b**) ligand-receptor (LR) pairs in PAD. In each panel, left part indicates the ligand log fold change (LFC) in *ATF3/4*^+^ ECs and right part the receptor LFC in *LYVE1*^hi^MHCII^low^ macrophages from each LR pair. Color indicates the LFC. **c**, Heatmap showing the regulatory potential of *ATF3/4*^+^ ECs ligands upregulated in PAD (left labels) over the downstream target genes upregulated in *LYVE1*^hi^MHCII^low^ macrophages (top labels). Color indicates the regulatory potential. **d**, Dot plot showing ORA results from the predicted targets in panel c. Color and size of the dots indicate -log10 adjusted p-value and number of genes in each category (intersection size), respectively. **e**, Graphs showing the geomean expression ± SEM of the markers CD206, CD80 and HLA-DR on human CD14^+^ cells cultured *in vitro* with increasing concentrations of ECs-derived conditioned media (CM). Non-ischemic CM (grey, n=3) and PAD CM (blue, n=5). Fisher’s one-tailed test (as implemented in g:Profiler) was used in d. Two-way ANOVA (* p-value < 0.05, *** p-value < 0.001) was used in e. P = 0.0002 (**e**, CD206), P = 0.0001 (**e**, CD80), P = 0.0414 (**e**, HLA-DR).

To explore the ability of ECs to influence *in vitro* monocyte to macrophage differentiation in PAD, we generated conditioned media (CM) from primary muscle ECs from non-ischemic (n=3) and PAD patients (n=5) and exposed it to CD14^+^ blood cells from healthy donors (n=6). FACS analysis showed that PAD-derived CM reduced CD206 expression (M2/regenerative) while increasing CD80 and HLA-DR (M1/inflammatory) in CD14^+^ cells (Fig. 7e), indicating that PAD EC-derived factors skew CD14^+^ cells toward a pro-inflammatory profile. Although these experiments used blood CD14^+^ cells rather than specific subtypes from our scRNAseq dataset, they provide supportive evidence that altered crosstalk in muscle ECs during PAD could influence immune cell behavior.

## Discussion

We generated a comprehensive scRNAseq dataset with over 100’000 cells from the muscle microenvironment of patients with PAD as well as non-ischemic co-morbidity matched controls (undergoing bypass surgery for an aneurysm). Pseudobulk analysis revealed increased TNFA and IL6 signaling, both associated with walking impairments in PAD patients^65,66^. Other upregulated processes such as inflammation, unfolded protein response and metabolism were also reported in previously bulk RNAseq studies of PAD muscle^67,68^. In contrast to our approach, those bulk studies are dominated by myonuclear transcripts, suggesting common pathway alterations across cell types. Our dataset provides a resource for cell-type-specific analysis of transcriptomics changes in PAD. While here we focused on specific cell types, other cell (sub)populations may also critically contribute to PAD progression.

Besides identifying common EC subpopulations, we also found *ATF3/4*^+^ venous and capillary ECs in human skeletal muscle, confirming earlier work in mice which showed that ATF3/4^+^ ECs are functionally required for exercise-induced angiogenesis^24^. A venous population, transcriptionally equivalent to *ATF3/4*^+^ venous, is specifically enriched in human skeletal muscle, adipose tissue and lymph nodes^69^. Also, glucosamine-driven stimulation of ATF4 signalling improves perfusion recovery in preclinical models of PAD^39^. Using this preclinical model, we showed that ATF4 deletion in ECs prevents ischemia recovery, further underscoring the critical contribution of ATF4 in ECs. Our scRNAseq and spatial data revealed fewer *ATF3/4*^+^ ECs in PAD, while also reducing their angiogenic and immune regulatory pathways. ECs in PAD upregulated genes related to EndoMT and cellular responses to metal ions. Elevated levels of these ions have been described in patients with PAD and atherosclerotic lesions^70–73^, and have been linked to endothelial dysfunction^74,75^. These results suggest that ECs undergo maladaptive adaptations that impair their functions in PAD. Through *in-silico* drug repurposing, we identified celastrol as a promising candidate to reprogram PAD ECs toward a healthier phenotype which highlights the potential of our scRNAseq data for therapeutic discovery.

Blood vessel formation and remodeling is heavily influenced by resident and recruited cells. In experimental models of PAD, monocyte-derived macrophages play a role in vascularization^16,19,21,40–42^ and muscle repair. After acute muscle ischemia or injury, macrophages adopt a pro-inflammatory/M1-like phenotype. Rapidly thereafter, M1 macrophages repolarize to anti-inflammatory (regenerative)/M2-like phenotype^76,77^, which is required for optimal muscle regeneration^77–79^. M2-like macrophages also have an important pro-angiogenic role since they are a source of VEGF. Nonetheless, acute mouse models differ from chronic PAD, so the contribution of macrophages in PAD remains unclear. We measured an almost 2-fold increase in CD68^+^ cells, predominantly *LYVE1*^hi^ macrophages and not monocyte-derived *LYVE1*^low^ macrophages. This observation is interesting since preclinical mouse models for muscle ischemia have unequivocally supported the requirement of monocyte-derived macrophages for muscle regeneration^80,81^. It is unclear whether the failed accumulation of these cells in PAD patients is a result of the chronic nature of PAD or contributing to the impaired regenerative response.

*LYVE1*^hi^MHCII^low^ macrophages were the dominant muscle macrophage population, even under a setting of chronic inflammation such as PAD. These cells were previously identified as embryonically-derived, locally self-renewing tissue resident macrophages^44,82^. They cloak microlesions to prevent widespread neutrophil recruitment^83^ or clear damage-induced apoptotic cells^47^, but their contribution to PAD muscle pathology is not known. Consistent with mouse literature, *LYVE1*^hi^MHCII^low^ macrophages in our human muscle samples expressed tissue-resident markers including *FOLR2* and other M2/regenerative marker genes such as *CD163* and *MRC1* (CD206). However, during PAD, *LYVE1*^hi^MHCII^low^ macrophages also activated a pro-inflammatory gene program. The co-expression of genes that are typically associated with an M1 as well as M2-like state might explain the contrasting findings reported in literature. Recently, TIMD4 was proposed as a marker for resident mouse muscle macrophages, but we could not detect *TIMD4* expression in human macrophages. Also, we found that *LYVE1*^hi^MHCII^low^ macrophages preferentially localize close to *ATF3/4*^+^ ECs in PAD and showed altered (more pro-inflammatory) crosstalk with ECs. Whether these altered EC-macrophage communication drives PAD muscle pathology or reflects a failed homeostatic response still needs further research.

Analysis of the transcriptomic changes in microvascular capillary ECs revealed partial EndoMT in PAD. EndoMT has been observed in atherosclerotic^84^ and PAD patients^5^, and ischemia promotes the activation of EndoMT in several pathological settings^85^. Also, partial EndoMT is required for optimal angiogenesis in myocardial infarction^32^. It is possible that persistent hypoxia contributed to a partial, and chronic EndoMT. While we did observe activation of some, but not all, markers of EndoMT in PAD capillary ECs, ECM genes were significantly activated. Also, the endothelial marker *CDH5* was reduced, but not lost, in PAD. Interestingly, this state of partial EndoMT might be reversible^32^.

We detected strong cellular crosstalk between different cell types that was significantly altered in PAD, with downregulated signaling networks related to muscle regeneration, angiogenesis and vessel junctions and upregulated pro-inflammatory pathways such as *IL1B* or *IL6*. IL1B is a known regulator of EndoMT^86^, and IL1B inhibitors improved maximum and pain-free walking distance in patients with symptomatic PAD^87^, although muscle pathology was not further investigated in this study. Analyzing SMCs/Pericytes-EC interactions revealed a reduction in *PGDFRB* and *NOTCH3* signaling in PAD, which control mural cells recruitment and attachment to the endothelium. Also, mural cells increased *NAMPT* signaling, which is associated with pro-inflammatory markers in CAD patients^59^. Both SMCs and pericytes upregulated contractility genes such as *ACTA2, TAGLN, MYH11* and *CNN1*, in line with increased contraction of vessels by pericytes under ischemia^88,89^.This phenotype might contribute to reduced perfusion in pathologies such as PAD^90^. Finally, we found that LYVE1^hi^MHCII^low^ macrophage activation could be EC-dependent. During PAD, ECs reduced *CSF1* and *GAS6* signaling with LYVE1^hi^MHCII^low^ macrophages. CSF1 is the primary growth factor responsible for macrophage differentiation and survival^91^, and endothelial CSF1 contributes to this^92^. Inhibition of GAS6 signaling in macrophages increases their inflammatory responses^93^, which would be consistent with the activated state of *LYVE1*^hi^MHCII^low^ macrophages and their increased cytokine production (Extended Data Fig. 5g). The EC ligands upregulated in PAD furthermore seem to control downstream genes in *LYVE1*^hi^MHCII^low^ macrophages related to inflammation, migration and cytokine response. We experimentally showed that secreted factors from ECs during PAD increased the expression of M1 markers while reducing M2 markers in human CD14^+^ immune cells, further supporting a more pro-inflammatory profile of ECs during PAD. Beyond having a perivascular localization, these data imply that ECs might control the fate of *LYVE1*^hi^MHCII^low^ macrophages, an exciting hypothesis that requires further investigation.

Our study has limitations. We included a non-ischemic control group with similar demographic and clinical risk factors as PAD patients to ensure that we detect PAD-specific differences rather than co-morbidity related effects. Patients with aneurysms have a similar clinical background of hypertension and hyperlipidemia, and a similar treatment of statins, anti-platelet therapy and antihypertensive drugs (Table 1). The non-ischemic patients had a normal ABI, did not show claudication symptoms during the walking test, therefore excluding the clinical manifestation of PAD, and did not show occluded vessels as reported by GLASS Stage. Additionally, popliteal aneurysm patients underwent a similar surgical procedure, which allowed us to standardize sample harvesting. We are confident that this strategy dissected differences between both groups as evidenced by the enrichment of gene sets related to abnormal muscle physiology. This said, we cannot exclude that vascular aneurysms affected the composition and genetic fingerprint of the non-ischemic samples. Our patient cohort was relatively small and, therefore, may be subjected to some inadvertent selection bias. Our study only included male participants, so potential sex-based differences could not be assessed.

In conclusion, we used scRNAseq to characterize the composition and transcriptional changes of the muscle microenvironment in ischemic PAD compared to non-ischemic co-morbidity matched patients. We identified profound transcriptomic alterations in ECs promoting EC dysfunction features (such as EndoMT) while acquiring a more pro-inflammatory phenotype. *LYVE1*^hi^MHCII^low^ macrophages are the main macrophage population in ischemic PAD muscle and get activated during the disease, a phenotype that may be driven by ECs. We propose that unravelling the impaired cellular crosstalk in PAD muscle could potentially offer novel targets for therapeutic intervention.

## Methods

### Patients and study design

Non-ischemic and PAD patients were recruited from the Department of Angiology, Cantonal Hospital Baden, Baden, Switzerland. The general **inclusion criteria for both non-ischemic and PAD patients** were: scheduled for surgery (bypass surgery in PAD patients or lower limb aneurysm surgery in non-ischemic samples); Age ≥45 years; ability to understand German; signed informed consent to participate in the study and received no compensation. **Inclusion criteria specific for PAD patients**: diagnosis of PAD ranging from exercised limited claudication (Fontaine classification: IIa or IIb) to CLI (Fontaine III-IV); ankle-brachial index (ABI) ≤ 0.90 at rest unless diagnosed with media sclerosis; stable medicament regimen (i.e., statin, anti-platelet, anti-hypertensive regimen). **Inclusion criteria specific for non-ischemic patients:** diagnosis lower limb aneurysm; ABI > 1 at rest. The general **exclusion criteria for both non-ischemic and PAD patients** were: severe peripheral neuropathy (Total Neuropathy Score ≥3); unstable angina or severe (>70%) three coronary vessel disease; significant liver dysfunction (Model for End-Stage Liver Disease Score) or known liver cirrhosis (Child-Turcotte-Pugh score >6 points); treated for cancer in the last 2 years; diagnosis of Parkinson’s disease. **Exclusion criteria specific for PAD patients:** above/below-knee amputation. **Exclusion criteria specific for non-ischemic patients:** peripheral artery disease diagnosed as described before (inclusion criteria specific for PAD patients). GLASS Stage: preoperative angiographic imaging (iodine contrast CT angiography or digital subtraction angiography) was performed. Then, for GLASS Stage calculation, the target arterial path (TAP) was identified and the femoropopliteal (FP) GLASS grade (NA, I, II, III) and the infrapopliteal (IP) GLASS grade (NA, I, II, III) were determined. FP and IP scores were combined to determine the final GLASS Stage. The study was conducted according to the Declaration of Helsinki, the Human Research Act (HRA) and the Human Research Ordinance (HRO) and the protocol was approved by the Ethics Committee of the Canton of Zurich (KEK number: 2020-01393). Remaining sample material is available upon formal request to the corresponding author.

Muscle samples from the peripheral region of the medial head of gastrocnemius muscle (1/3 of the distance between the tibial plateau and the malleolus medialis) were collected during lower limb aneurysm surgery (non-ischemic samples) or bypass surgery (PAD samples) and immediately transferred to an ice-cold hypothermic preservation solution (HypoThermosol® FRS Preservation Solution, H4416 Sigma-Aldrich) for optimal transport and conservation until further processing. From 4 PAD and 4 non-ischemic samples, samples were processed for scRNAseq analysis, while part of the tissue was also used for histology. A set of samples (n= 11 for PAD; n= 8 for non-ischemic) was embedded in optimal cutting temperature medium, frozen in liquid nitrogen-cooled isopentane and stored at -80°C for histological analysis. Transversal 10-μm cryosections of the samples were prepared at -20°C, air dried for ∼30 min and stored at -80°C until further processing. A subset of histology samples (n = 3 for PAD; n = 3 for non-ischemic) was used for CosMx single-cell spatial transcriptomics (see below). A set of 8 independent samples (3 non-ischemic vs 5 PAD) was used for isolation and culture of primary muscle ECs (see below).

### Isolation of cells from human skeletal muscle for single cell RNA sequencing

For isolation of cells for scRNAseq, and in less than 2 hours after collection and preservation in ice-cold hypothermic preservation solution (HypoThermosol® FRS Preservation Solution, H4416 Sigma-Aldrich), samples were minced in a petri dish on ice using scissors and a surgical blade. Minced samples were transferred into a digestion buffer (4 mL digestion buffer per 100 mg of muscle) containing 2 mg/ml Dispase II (D4693 Sigma-Aldrich) and 2 mg/ml Collagenase IV (17104019 Thermo Fisher Scientific) in HBSS buffer (14025100 Thermo Fisher Scientific) at 37°C for 25-30 minutes, with gentle shaking every 3-5 minutes. After digestion, the reaction was stopped by adding an equal volume of 20% heat inactivated FBS (10500064 Thermo Fisher Scientific) 1mM EDTA (E177 VWR Life Science) in HBSS (14025100 Thermo Fisher Scientific) and the suspension was passed through a 100-μm cell strainer (800100 Bioswisstec) and a 40-μm cell strainer (352340 Falcon). Cell suspension was centrifuged at 500 g for 5 minutes at 4°C and the supernatant discarded. The pellet was resuspended in 1 mL of ACK Lysing Buffer (A1049201 Thermo Fisher Scientific) for red blood cell lysis for 2 minutes at room temperature. After incubation, the reaction was stopped with 4 mL of 10% heat inactivated FBS (10500064 Thermo Fisher Scientific) in HBSS buffer (14025100 Thermo Fisher Scientific). Cell suspension was centrifuged at 500 g for 5 minutes at 4°C and the pellet resuspended in 0.5% BSA (A1391 PanReac AppliChem) in DPBS (14190250 Thermo Fisher Scientific) containing 1mM EDTA (E177 VWR Life Science) for FACS. Calcein Violet (65-0854-39 Thermo Fisher Scientific) for metabolically active live cells selection was added directly to the cell suspension at 1:1000 dilution 10 minutes before sorting.

### Human scRNAseq analysis

Metabolically active (Calcein^+^) human muscle mononuclear cells were FACS sorted and loaded into the 10X Chromium Next GEM Chip G, aiming the recovery of ∼10000 cells per sample. scRNAseq libraries were generated according to 10X Chromium Next Gem Single Cell 3’ Reagent Kits v3.1 (CG000204 Rev D) and sequenced on the Illumina Novaseq 6000 system. We sequenced a total of 13 samples from 8 different patients (Table 1). Sequencing reads were aligned to the human genome using CellRanger v5.0.0-7.0.0 and the genome assembly GENCODE GRCh38.p13 Annotation Release 37. Intron reads were not included during the alignment. Each individual sample was independently processed for quality control (QC) and log-normalization using *scran* 1.24.1 and scater 1.24.0 packages in R (version 4.2.0). High quality cells were selected based on library size, number of detected genes and percentage of reads mapped to mitochondrial genes, independently adjusting the thresholds for each sample. Raw UMI counts were log-normalized based on library size factors. Doublets were detected and removed in every individual sample using the default parameters of the function *runDoubletFinder* from *singleCellTK* 2.6.0 package. After individual QC and log-normalization, all samples were batch-corrected and merged using *Harmony* 0.1.1 package. Dimensionality reduction by t-Distributed Stochastic Neighbor Embedding (TSNE) was performed based on the dimensions generated by *Harmony* integration. Clustering was performed by building a nearest neighbor graph (*k* = 20) and then applying the Louvain algorithm as implemented in *Seurat* 4.1.1 package. For individual analysis and reclustering of the EC, monocyte/macrophage and SMC/Pericytes populations, cells were selected by tracing back the clustering of the complete dataset in every individual sample. Raw counts from selected cells, either ECs, monocyte/macrophage and SMC/Pericytes were log-normalized in every individual sample and batch-corrected and merged as described above for the complete dataset. All dotplots showing gene expression were generated using the *plotDots()* function as implemented in scater 1.24.0 package. Cell type proportions in each condition were calculated by dividing the number of cells in each cluster by the total number of cells in that respective condition.

Differentially expressed genes (DEGs) were detected by performing a Wilcoxon Rank Sum test as implemented in *Seurat* 4.1.1 package. DEGs were selected based on a minimum log_2_ fold change of 0.25 and an adjusted p-value < 0.05. For over-representation analysis (ORA), selected DEGs were ordered based on increasing adjusted p-value and given as an input in the web-based implementation of g:Profiler^26^, selecting the option “Ordered query” before running the analysis. ORA results were represented as the -log10 of the p-values obtained for each process. ORA analysis results were simplified by using *rrvgo* 1.10.0 package. Redundant terms were grouped and only the most significant term of each group was conserved. Schematic visualization of KEGG pathways from ORA results were generated using *pathview* 1.38.0 package.

For pseudobulk analysis, lowly expressed genes were removed if they were not detected in at least 10 cells. Raw UMI counts were aggregated per patient (total sum of counts for every gene from all cells in every patient). For differential expression analysis in the pseudobulk dataset, we applied *DESeq2* 1.38.3 package and performed the comparison between non-ischemic and PAD patients. All significant DEGs (adjusted p-value <0.05) were reported in a heatmap (Fig. 1j) by using the normalized counts (by size factors) in each patient and scaled and centered based on gene expression per patient. For Gene Set Enrichment Analysis (GSEA) in the pseudobulk dataset we applied *fgsea* 1.24.0 package. From the differential expression analysis results performed with DESeq2 we selected the *stat* value (Wald statistic: the log2FoldChange divided by lfcSE) for each gene and performed GSEA analysis using the default settings of *fgsea()* function from *fgsea* 1.24.0 package. Significant pathways were selected based on an adjusted p-value < 0.05. Human Hallmark pathways from MsigDB^94^ were used for this analysis.

Gene Set Variation Analysis was performed using *GSVA* 1.46.0 package. Log-normalized counts from the corresponding scRNAseq dataset (either ECs or Monocytes/Macrophages) were given as an input for the *gsva()* function as implemented in GSVA 1.46.0 package. The gene sets to perform GSVA were obtained from Fan et al^24^ (ATF4 dependent genes as defined by genes with logFC < -0.1 and adjusted p-value < 0.05 after comparing mouse muscle ECs in wild-type and EC-specific inducible ATF4 knockout mice) or Sanin et al^48^ (macrophage activation processes). For GSVA of M1-M2 signatures, we obtained the genesets from Varga et al^50^, which reported a set of M1-M2 genes derived from *in vitro* stimulation and generated genesets for pro-inflammatory (Ly6C^+^) and pro-regenerative (Ly6C^-^) macrophages at different timepoints after cardiotoxin-induced muscle injury.

Bulk RNAseq data from muscle ECs in WT and EC-specific ATF4 KO mice were obtained from Fan et al^24^ and processed as described by the authors. Only WT and ATF4 KO red muscle ECs (RmECs) were used in this study.

Transcription factor inference analysis was performed using the Python implementation of SCENIC package (pySCENIC)^30^ and following the script provided by the authors in their GitHub repository (https://github.com/aertslab/pySCENIC). Cell communication analysis on the complete dataset (all cell types in non-ischemic and PAD samples) was performed using *CellChat* 1.6.1 package^53^ following the script provided by the authors in their GitHub repository (*Full tutorial for comparison analysis of multiple datasets* section in https://github.com/sqjin/CellChat). Cell communication analysis between *ATF3/4*^+^ ECs and *LYVE1*^hi^MHCII^low^ macrophages was performed using *nichenetr* 1.1.0 package^64^ following the script provided by the authors in their GitHub repository (*Differential NicheNet analysis between conditions of interest* section in https://github.com/saeyslab/nichenetr).

All raw and processed (10X CellRanger output files) sequencing data are available at GSE235143.

### Isolation and culture of endothelial cells from human skeletal muscle

We isolated muscle ECs from 8 independent samples (3 non-ischemic vs 5 PAD) using an established protocol^95^, yielding high-purity (>95%) EC cultures (Extended Data Fig. 4g). As recommended in the original publication^95^, we performed several rounds of CD31^+^ magnetic beads sorting until we achieved the desired purity. Cells were cultured in EGM^TM^-2 MV Microvascular Endothelial Cell Growth Medium-2 BulletKit^TM^ (CC-3202, Lonza). Subsequently, confluent EC cultures from 3 PAD patients were treated for 24h with vehicle (DMSO, A994.1, Carl Roth) or celastrol (C0869, Sigma-Aldrich) (125 or 250 nM), doses that are consistent with literature and do not affect cell viability of HUVECs^38^. Celastrol powder was dissolved in DMSO at 2 mg/mL (stock concentration) and later diluted to experimental doses (125 or 250 nM) directly into EGM-2 media right before use. Control samples (Vehicle) were given the same volume of DMSO as used to dilute celastrol in the 250 nM condition. After 24h incubation, RNA was extracted for bulk RNA sequencing using RNeasy Plus Micro Kit (74034, Qiagen) according to the manufacturer’s protocol. RNA integrity was assessed by using Agilent High Sensitivity RNA ScreenTape System. Only samples with RNA Integrity Number (RIN) ≥ 8.0 were further processed.

For conditioned media experiments, confluent ECs were cultured for 48h with fresh EGM^TM^-2 Medium. Then, media was collected, filtered through a 0.2 μm filter (83.1826.001, Sarstedt) and cryopreserved at -80°C until further use. Cells were routinely cultured at 37°C in 21% O_2_ and 5% CO_2_.

### Macrophage polarization in vitro using conditioned media

*In vitro* macrophage polarization using conditioned media was performed with an adapted protocol from George et al^96^. PBMCs were obtained from buffy coats (Swiss Blood Donation Center of Lugano) by density gradient centrifugation using Ficoll® Paque Plus (17-1440-02, Cytiva). CD14^+^ cells were isolated using human CD14 MicroBeads (130-050-201, Miltenyi Biotec) following the manufacturer’s recommendations. The purity of CD14^+^ cells was assessed by flow cytometry (>98%, except from 1 donor, >90%) (see antibodies below). 5×10^5^ CD14^+^ cells per well were seeded in a 48-well plate (TPP92148, TPP) in 250 μl of varying concentrations (0%, 6.25%, 12.5%, 25%, 50%) of conditioned media (3 non-ischemic, 5 PAD). On day 3, the cells were harvested, stained for flow cytometry and acquired on a BD Fortessa. The following antibodies were used: Human TruStain FcX (used for Fc blocking; 422302, BioLegend), Zombie NIR™ Fixable Viability Kit (dilution 1:500, used for Live/Dead staining; 423106, BioLegend), CD14 (APC-Cy7, dilution 1:200, used for purity assessment; 325620, BioLegend), CD19 (FITC, dilution 1:20, used for purity assessment; 555412, BD Biosciences), CD3 (BV650, dilution 1:200, used for purity assessment; 300468, BioLegend), CD56 (PC5, dilution 1:50, used for purity assessment; A07789, Beckman Coulter), HLA-DR (BV605, dilution 1:100, used for polarization assessment; 562845, BD Biosciences), CD80 (FITC, dilution 1:50, used for polarization assessment; 557226, BD Biosciences), and CD206 (PE, dilution 1:50, used for polarization assessment; 321106, BioLegend).The data was analyzed using FlowJo v10. The experiment was performed twice with CD14^+^ cells from 3 buffy coats, respectively (total of 6 different donors).

### Bulk RNA sequencing of celastrol-treated ECs

Libraries were generated using TruSeq Stranded mRNA Library Kit (Illumina) and sequenced on the Illumina NovaSeq X plus. Sequencing reads were processed using Kallisto and the human genome assembly GENCODE GRCh38.p13 Annotation release 42 to generate a count file matrix for each individual sample. Samples were pooled together into a single matrix and analyzed following DESeq2 pipeline. Differential expression analysis was performed after correcting for patient and sequencing plate effect. Gene signatures for GSEA were obtained from Fan et al^24^ (ATF4, see “Human scRNAseq analysis” section) and Alhusban et al^39^ (Glucosamine, after analyzing bulk RNAseq data from original publication, HUVECs treated with glucosamine (5 mM) vs L-glucose (5 mM) for 6 hours under hypoxia and serum starvation (HSS) conditions).

Raw and processed (Kallisto aligned) sequencing files are available at GSE287300.

### CosMx single-cell spatial transcriptomics of muscle tissue

Transversal 10-μm cryosections of the samples were prepared at -20°C, air dried for ∼30 min and shipped to Nanostring (Seattle, WA, USA) under optimal storage conditions using dry ice to ensure sample integrity. Samples were processed by Nanostring for CosMx SMI through the Technology Access Program (TAP, project ID: SMI0364). For transcript detection, CosMx Human Universal Cell Characterization Panel (RNA, 1000 Plex; CMX-H-USCP-1KP-R) was used. Cell detection (cell segmentation) was performed using a combination of DAPI (nuclei) staining and laminin staining, which marks the basal lamina surrounding individual muscle fibers to provide additional structural context and facilitate accurate segmentation of mononuclear cells located between fibers. Data processing was performed using Giotto package. Each sample was individually processed for quality control and log-normalization and then integrated together using Harmony, following the instructions from Giotto developers (https://github.com/drieslab/Giotto). To integrate spatial samples with scRNAseq data and transfer cluster labels, spatial samples were treated as scRNAseq-like datasets (cell-by-gene expression matrices). These were integrated with scRNAseq samples using the same Harmony pipeline described in the “Human scRNAseq analysis” section. After integration, all cells were projected into a shared dimensionality reduction space. We then used TSNE coordinates to identify the 10 closest scRNAseq cells for each spatial cell, using Euclidean distance: 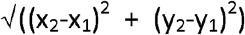. The most frequent cluster label among the 10 nearest scRNAseq cells was assigned to the corresponding spatial cell. For proximity analysis between LYVE1^hi^MHCII^low^ macrophages and EC subtypes a similar approach was followed. Briefly, after transferring cluster labels to the spatial dataset, we identified the nearest EC for each LYVE^hi^MHCII^low^ macrophage based on the shortest Euclidean distance in the spatial coordinates.

### Mice

Inducible EC-specific Atf4 knockout (*Pdgfb*-Cre^ERT2^ x *Atf4*^ΔEC/ΔEC^) mice were generated as described before^24^. Recombination was induced in 8-10 weeks old mice by daily intraperitoneal (IP) administration of 1 mg tamoxifen (T5648, Sigma-Aldrich) dissolved in 1:10 ethanol:corn oil solution for 5 consecutive days. A wash out period of at least 7 days was allowed before starting the experiments. Tamoxifen-treated Cre-negative littermates were used as control for all experiments. All mice were maintained on a C57BL/6N background. To label proliferating cells, an IP injection of 5-ethynyl-2’-deoxyuridine (EdU) (E10187, Thermo Fischer Scientific) solution (5 mg/ml in saline) was performed 7 hours before sacrificing the mice.

Mice were randomly allocated to different treatment groups, and the investigator was blinded to the group allocation during the experiment as well as during the analysis. Mice were housed at standard housing conditions (22°C, 12 h light/dark cycle), with ad libitum access to chow diet (18% proteins, 4.5% fibers, 4.5% fat, 6.3% ashes, Provimi Kliba SA) and water. Health status of all mice was regularly monitored according to FELASA guidelines. All animal experiments were approved by the local animal ethics committee (Kantonales Veterinärsamt Zürich, license ZH050/2021), and performed according to local guidelines (TschV, Zurich) and the Swiss animal protection law (TschG). All mice in this study were male.

### Hindlimb ischemia model

Hindlimb ischemia experiments were performed as described before^42^. Mice were anesthetized with isofluorane, the hindlimbs were shaved, and, following a small incision in the skin, both the proximal end of the femoral artery and the distal portion of the saphenous artery were ligated. The artery and all side-branches were dissected free; after this, the femoral artery and attached side-branches were excised. Immediately after surgery, perfusion was measured by Laser Doppler Imaging of plantar regions of interest (Moor Instruments Ltd, Axminster, Devon, England) and calculated as ratio of left (ligated) versus right (unligated) values. Perfusion was measured again at different timepoints following the same procedure.

### Tissue Immunofluorescence and Histology

Human muscles were harvested and embedded in OCT embedding matrix (6478.1, Carl Roth) and frozen in liquid N_2_-cooled isopentane. Frozen sections (10 μm) were made using a cryostat (Leica CM 1950). The following antibodies were used: anti-LYVE1 (1:200 dilution, ab33682, Abcam), anti-CD68 (1:100 dilution, ab955, Abcam), anti-CD31 (1:200 dilution, M082329-2, Dako), anti-Fibronectin (1:1000 dilution, ab23750, Abcam), Goat anti-Rabbit IgG Cross-Adsorbed Secondary Antibody, Alexa Fluor 568 (1:200 dilution, A-11001, Thermofisher), Goat anti-Mouse IgG Cross-Adsorbed Secondary Antibody, Alexa Fluor 488 (1:200 dilution, A-1101, Thermofisher), Alexa Fluor 647 conjugated wheat germ agglutinin (WGA, 1:50 dilution, W32466, ThermoFisher). Muscle sections were fixed in 4% paraformaldehyde (PFA) followed by epitope retrieval using sodium citrate (10 mM, pH 6.5) at 92 °C for 20 min. Endogenous peroxidase activity was blocked with 3% hydrogen peroxide in phosphate-buffered saline followed by 10 min of permeabilization (PBS + 0.5% Triton X-100) and 60 min of blocking at room temperature in blocking buffer (PBS + 1% BSA). After blocking, sections were stained overnight at 4°C with primary antibodies, followed by 1 hour incubation of secondary antibodies at room temperature. Extracellular matrix was stained using wheat germ agglutinin (WGA). Nuclei were detected using Hoechst diluted 1:5000 in PBS and for 5 minutes at room temperature. Slides were then washed with PBS and mounted using Immu-Mount (9990412, Thermofisher). Slides were imaged on an AxioObserver.Z1 fluorescence microscope using a 20x and 40x objective (Carl Zeiss, Oberkochen, Germany). For the LYVE1 and CD68 staining, 315 ± 106 (mean ± standard deviation [SD]) fibers were counted per patients, and for the CD31 staining, 409 ± 80 (mean ± SD) fibers were counted. Different regions over the whole tilescan muscle section were analyzed with the Zen software (ZEN 2011 imaging software, Zeiss) and Fiji. General tissue morphology was evaluated using haematoxylin and eosin (H&E) staining. H&E images were acquired with an Eclipse Ti2 inverted microscope (Nikon) using a 20x objective.

Mouse muscles were harvested and embedded in OCT embedding matrix (6478.1, Carl Roth) and frozen in liquid N_2_-cooled isopentane. Frozen sections (10 μm) were made using a cryostat (Leica CM 1950). For EdU detection combined with CD31, EdU was first visualized using the EdU Click-iT Cell Reaction Buffer Kit (C10269, ThermoFisher) according to manufacturer’s instructions, and subsequently incubated for 1h in blocking buffer (PBS with 1% BSA) at room temperature. Thereafter, sections were incubated overnight at 4°C with goat anti-Mouse/Rat CD31/PECAM-1 antibody (1:250, 3628, R&D Systems) diluted in blocking buffer with 0.1% Triton X-100.

### Statistics

The histology images presented in the manuscript are representative of the data (quantification of images is approximately the group average) and the staining quality. All human imaging data are represented as violin plots, where the three horizontal lines represents the median and quartiles, and the width of the plot indicates the distribution of the data. GraphPad Prism software (version 9.2.0) was used for statistical analyses. Shapiro-Wilk test was performed to analyze data distribution. Non-normally distributed data were analyzed by non-parametric Mann-Whitney U test to compare two groups. Normally distributed data were analyzed using Student’s t-test in an unpaired two-tailed fashion to compare two groups. For experiments evaluating more than one variable, a two-way ANOVA with Tukey’s multiple comparison was used. For experiments evaluating more than one variable with a repeated measures design, a two-way ANOVA with Šidák’s multiple comparison test was used. Each figure legend indicates the statistical approach for each experiment displayed in the figure. p > 0.05 is considered non-significant (ns). Asterisks in figures denote statistical significance. R and related packages were used for RNA-seq statistical analysis. Unless otherwise indicated, the default statistical test (e.g., Wilcoxon Rank Sum test for differential expression) from each package function was used. No animals or data points were excluded from the analysis.

## Data availability

The RNA sequencing data underlying this article can be fully explored at https://shiny.debocklab.hest.ethz.ch/Turiel-et-al/. Additionally, raw data are available in Gene Expression Omnibus (GEO) at https://www.ncbi.nlm.nih.gov/geo/ and can be accessed with accession number GSE235143 (scRNAseq) and GSE287300 (celastrol bulk RNAseq). Processed data (scRNAseq datasets after QC, normalization, integration and clustering) can be accessed at Figshare^97^ (https://doi.org/10.6084/m9.figshare.29493215).

## Code availability

The following codes were used: SCENIC (https://github.com/aertslab/pySCENIC), CellChat (https://github.com/sqjin/CellChat), NicheNet (https://github.com/saeyslab/nichenetr) and Giotto (https://github.com/drieslab/Giotto). Any additional code required to reproduce the results from this manuscript is available from the corresponding author upon request.

## Acknowledgments

The authors thank the digital trial intervention platform (dTIP) from ETH Zurich for their skilled assistance, the Functional Genomics Center Zürich (FGCZ) and the Scientific Center for Optical and Electron Microscopy (ScopeM) for excellent technical support.

This work was supported by the Swiss National Science Foundation (SNSF) [grant number 310030_208041] and Vontobel Foundation [grant number 0521/2023]. K.D.B. is endowed by the Schulthess Foundation. D.Latorre is supported by the Helmut Horten Foundation, the Novartis Foundation for medical-biological Research and the GBS/CIDP Foundation International (ETH Zurich), and the Giovanni Armenise Harvard Foundation Career Development Award (OSR, Milan).

## Contributions

G.T. conceptualized the study, designed and performed the experiments and bioinformatic analyses, and wrote the paper. T.D. performed human sample processing and FACS sorting for scRNAseq. E.M. and D.Lussi performed and analyzed the histological experiments. Z.F. performed and analyzed the mouse experiments. C.M.C and D.Latorre performed and supervised the conditioned media experiments. G.B. performed the GLASS scoring. R.A., K.S., M.B. and J.Z. contributed to data analysis, interpretation, and the performance of experiments. S.F.F provided human muscle samples. S.E. conceptualized the study, recruited the patients, and performed the surgeries for human sample collection. K.D.B. conceptualized the study, supervised the experiments, acquired funding, and wrote the paper. All authors approved the final version of the paper.

