## Extended Data for "Single cell compendium of muscle microenvironment in peripheral artery disease reveals altered endothelial diversity and LYVE1^+^ Macrophage Activation"

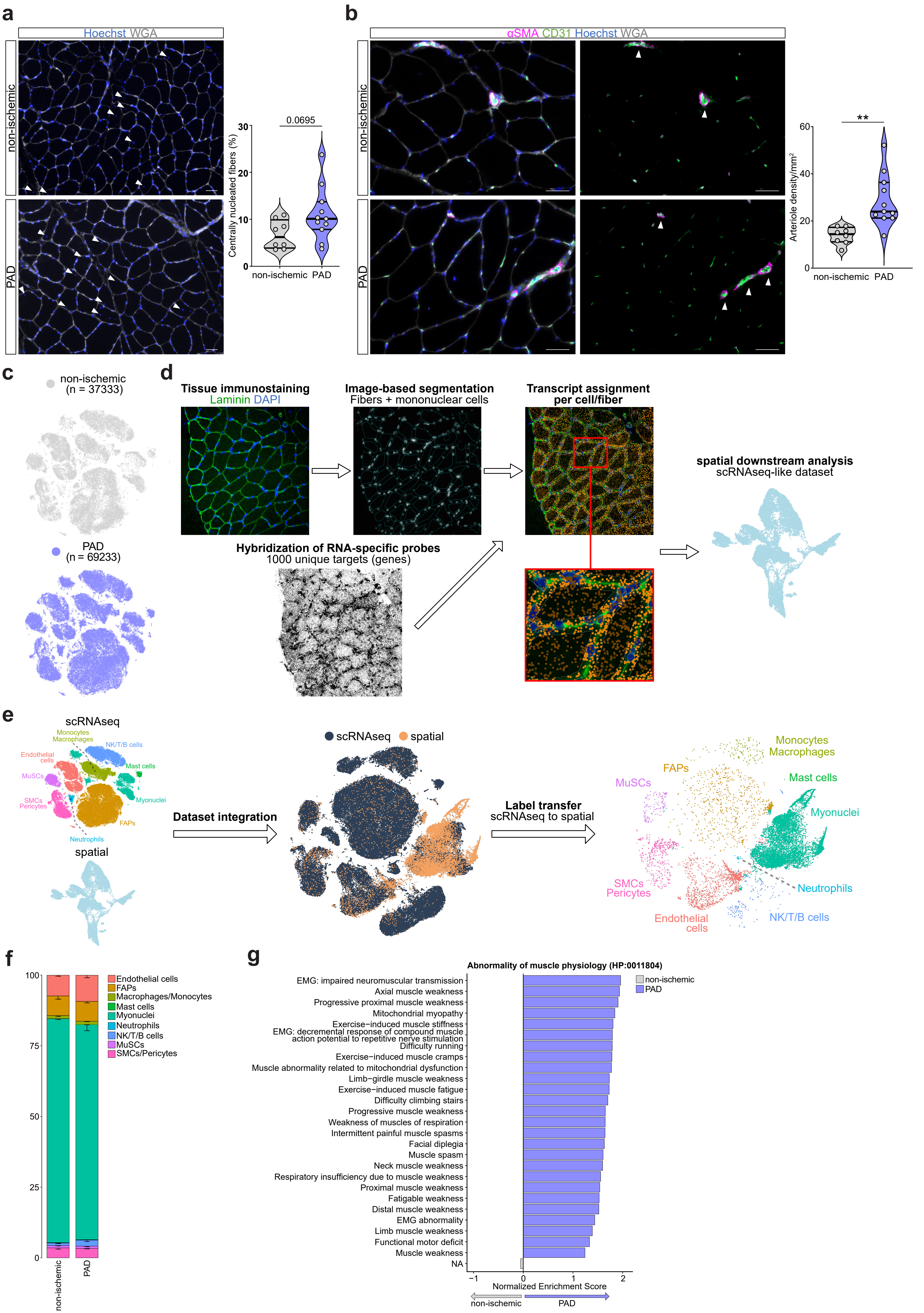

a
Hoechst WGA
non-ischemic
PAD
0.0695
Centrally nucleated fibers (%)
30
20
10
0
non-ischemic PAD
b
αSMA CD31 Hoechst WGA
non-ischemic
PAD
**
Arteriole density/mm²
60
40
20
0
non-ischemic PAD
c
non-ischemic
(n = 37333)
PAD
(n = 69233)
d
Tissue immunostaining
Laminin DAPI
Image-based segmentation
Fibers + mononuclear cells
Transcript assignment
per cell/fiber
Hybridization of RNA-specific probes
1000 unique targets (genes)
spatial downstream analysis
scRNAseq-like dataset
e
scRNAseq
Monocytes
Macrophages
NK/T/B cells
Endothelial cells
Mast cells
MuSCs
SMCs
Pericytes
Myonuclei
FAPs
Neutrophils
spatial
Dataset integration
scRNAseq spatial
Label transfer
scRNAseq to spatial
Monocytes
Macrophages
FAPs
MuSCs
Mast cells
Myonuclei
SMCs
Pericytes
Neutrophils
Endothelial cells
NK/T/B cells
f
100
75
50
25
0
non-ischemic
PAD
Endothelial cells
FAPs
Macrophages/Monocytes
Mast cells
Myonuclei
Neutrophils
NK/T/B cells
MuSCs
SMCs/Pericytes
g
Abnormality of muscle physiology (HP:0011804)
non-ischemic
PAD
EMG: impaired neuromuscular transmission
Axial muscle weakness
Progressive proximal muscle weakness
Mitochondrial myopathy
Exercise−induced muscle stiffness
EMG: decremental response of compound muscle action potential to repetitive nerve stimulation
Difficulty running
Exercise−induced muscle cramps
Muscle abnormality related to mitochondrial dysfunction
Limb−girdle muscle weakness
Exercise−induced muscle fatigue
Difficulty climbing stairs
Progressive muscle weakness
Weakness of muscles of respiration
Intermittent painful muscle spasms
Facial diplegia
Muscle spasm
Neck muscle weakness
Respiratory insufficiency due to muscle weakness
Proximal muscle weakness
Fatigable weakness
Distal muscle weakness
EMG abnormality
Limb muscle weakness
Functional motor deficit
Muscle weakness
NA
−1
0
1
2
Normalized Enrichment Score
non-ischemic
PAD

**Extended Data Fig. 1: scRNAseq reveals cell heterogeneity in PAD. a,** Representative images of gastrocnemius muscle from non-ischemic (n=8) and PAD (n=11) stained for nuclei (Hoechst, blue) and WGA (white) (Scale bar: 50 µm) and quantification of percentage of centrally nucleated fibers over total fibers. **b,** Representative images of gastrocnemius muscle from non-ischemic (n=8) and PAD (n=11) stained for αSMA (red), CD31 (green), nuclei (Hoechst, blue) and WGA (white) (Scale bar: 50 µm) and quantification of number of arterioles per $mm^2$. **c,** TSNE plot (Fig. 1g) separated by condition, color indicates the condition. **d,** Schematic representation of the pipeline used for generating the spatial transcriptomics dataset (see Methods). **e,** Schematic representation of the pipeline used for transferring cluster labels from scRNAseq to spatial data (see Methods). **f,** Stacked bar plots showing cluster percentage in each condition in the spatial dataset from non-ischemic (n=3) and PAD (n=3) patients, color-coded by cluster. Each stack represents mean - SEM. **g,** Bar plots showing Normalized Enrichment Score of significant (adjusted p-value < 0.05) “Abnormality of muscle physiology” pathways (Human Phenotype Ontology) from GSEA analysis on the pseudobulk dataset, color indicates the condition. Each dot represents a single patient in panels a and b. Student’s t test (two-tailed, unpaired, parametric, ns > 0.05) was used in a and b. Adaptive multi-level Monte-Carlo scheme (as implemented in fgsea package) was used in g. P = 0.0695 (**a**), P = 0.0019 (**b**).

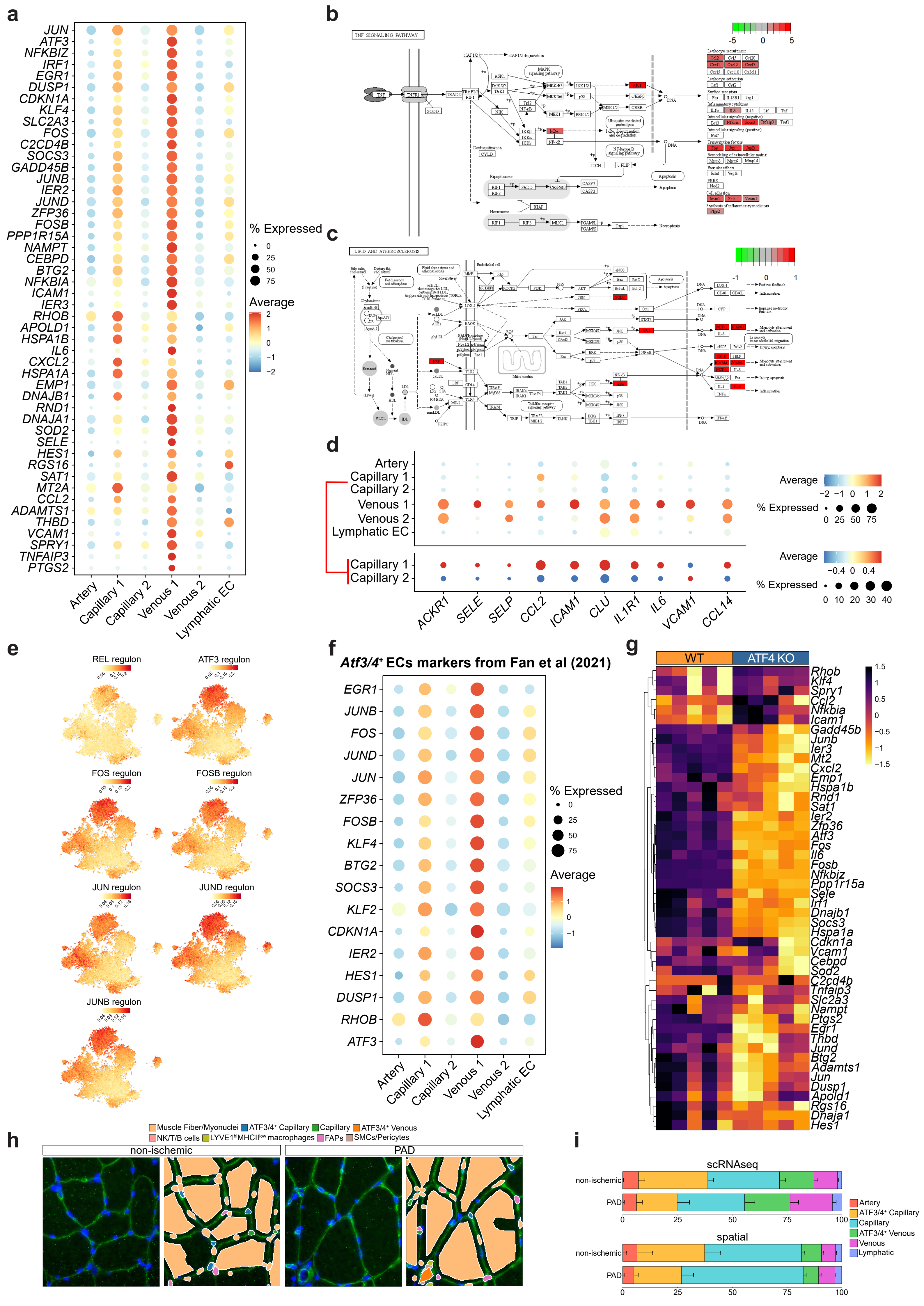
a
JUN
ATF3
NFKBIZ
IRF1
EGR1
DUSP1
CDKN1A
KLF4
SLC2A3
FOS
C2CD4B
SOCS3
GADD45B
JUNB
IER2
JUND
ZFP36
FOSB
PPP1R15A
NAMPT
CEBPD
BTG2
NFKBIA
ICAM1
IER3
RHOB
APOLD1
HSPA1B
IL6
CXCL2
HSPA1A
EMP1
DNAJB1
RND1
DNAJA1
SOD2
SELE
HES1
RGS16
SAT1
MT2A
CCL2
ADAMTS1
THBD
VCAM1
SPRY1
TNFAIP3
PTGS2
Artery
Capillary 1
Capillary 2
Venous 1
Venous 2
Lymphatic EC
% Expressed
0
25
50
75
Average
2
1
0
−1
−2
b
TNF SIGNALING PATHWAY
-5
0
5
c
LIPID AND ATHEROSCLEROSIS
-1
0
1
d
Artery
Capillary 1
Capillary 2
Venous 1
Venous 2
Lymphatic EC
Capillary 1
Capillary 2
ACKR1
SELE
SELP
CCL2
ICAM1
CLU
IL1R1
IL6
VCAM1
CCL14
Average
−2
−1
0
1
2
% Expressed
0
25
50
75
Average
−0.4
0
0.4
% Expressed
0
10
20
30
40
e
REL regulon
ATF3 regulon
FOS regulon
FOSB regulon
JUN regulon
JUND regulon
JUNB regulon
f
Atf3/4+ ECs markers from Fan et al (2021)
EGR1
JUNB
FOS
JUND
JUN
ZFP36
FOSB
KLF4
BTG2
SOCS3
KLF2
CDKN1A
IER2
HES1
DUSP1
RHOB
ATF3
Artery
Capillary 1
Capillary 2
Venous 1
Venous 2
Lymphatic EC
% Expressed
0
25
50
75
Average
1
0
−1
g
WT
ATF4 KO
Rhob
Klf4
Spry1
Ccl2
Nfkbia
Icam1
Gadd45b
Junb
Ier3
Mt2
Cxcl2
Emp1
Hspa1b
Rnd1
Sat1
Ier2
Zfp36
Atf3
Fos
Il6
Fosb
Nfkbiz
Ppp1r15a
Sele
Irf1
Dnajb1
Socs3
Hspa1a
Cdkn1a
Vcam1
Cebpd
Sod2
C2cd4b
Tnfaip3
Slc2a3
Nampt
Ptgs2
Egr1
Thbd
Jund
Btg2
Adamts1
Jun
Dusp1
Apold1
Rgs16
Dnaja1
Hes1
1.5
1
0.5
0
−0.5
−1
−1.5
h
Muscle Fiber/Myonuclei
ATF3/4+ Capillary
Capillary
ATF3/4+ Venous
NK/T/B cells
LYVE1hiMHCIIlow macrophages
FAPs
SMCs/Pericytes
non-ischemic
PAD
i
scRNAseq
non-ischemic
PAD
0
25
50
75
100
spatial
non-ischemic
PAD
0
25
50
75
100
Artery
ATF3/4+ Capillary
Capillary
ATF3/4+ Venous
Venous
Lymphatic

**Extended Data Fig. 2: A subpopulation of ECs shows an *ATF3/4* transcriptomic signature. a,** Dot plot of centered logcounts values from DEGs (adjusted p-value < 0.05 and log2FC > 1) upregulated in Venous 1 compared to Venous 2. **b-c**, Schematic view of TNF signaling (**b**) and lipid and atherosclerosis (**c**) pathways from KEGG (see Fig. 2d). Color indicates the log2FC of genes upregulated in Venous 1 compared to Venous 2. **d,** Dot plot of centered logcounts values from immunomodulatory ECs (IMECs) markers obtained from Alnaqbi et al. **e**, TSNE plots showing the SCENIC regulon activity score for REL and ATF4-dependent transcription factors. Color indicates the regulon activity score. **f,** Dot plot of centered logcounts values from mouse *ATF3/4*$^{+}$ Capillary marker genes obtained from Fan et al. **g,** Heatmap of centered normalized values of genes from panel a in the bulk RNAseq experiment of muscle ECs between WT and EC-specific ATF4 KO mice from Fan et al. Color indicates the centered normalized value. **h,** Images of non-ischemic and PAD muscle samples annotated in the spatial dataset after transferring cell labels from the scRNAseq dataset (see Extended Data Fig. 1e). Stained for laminin (green) and cell nuclei (blue, DAPI). Legend only shows visible cell types in the images. **i,** Stacked bar plots showing cluster percentage in each condition in the scRNAseq (top, non-ischemic n=4, PAD n=4) and spatial (bottom, non-ischemic n=3, PAD n=3) datasets, color-coded by cluster. Each stack represents mean - SEM. Wilcoxon Rank Sum test (as implemented in Seurat package) was used in a.

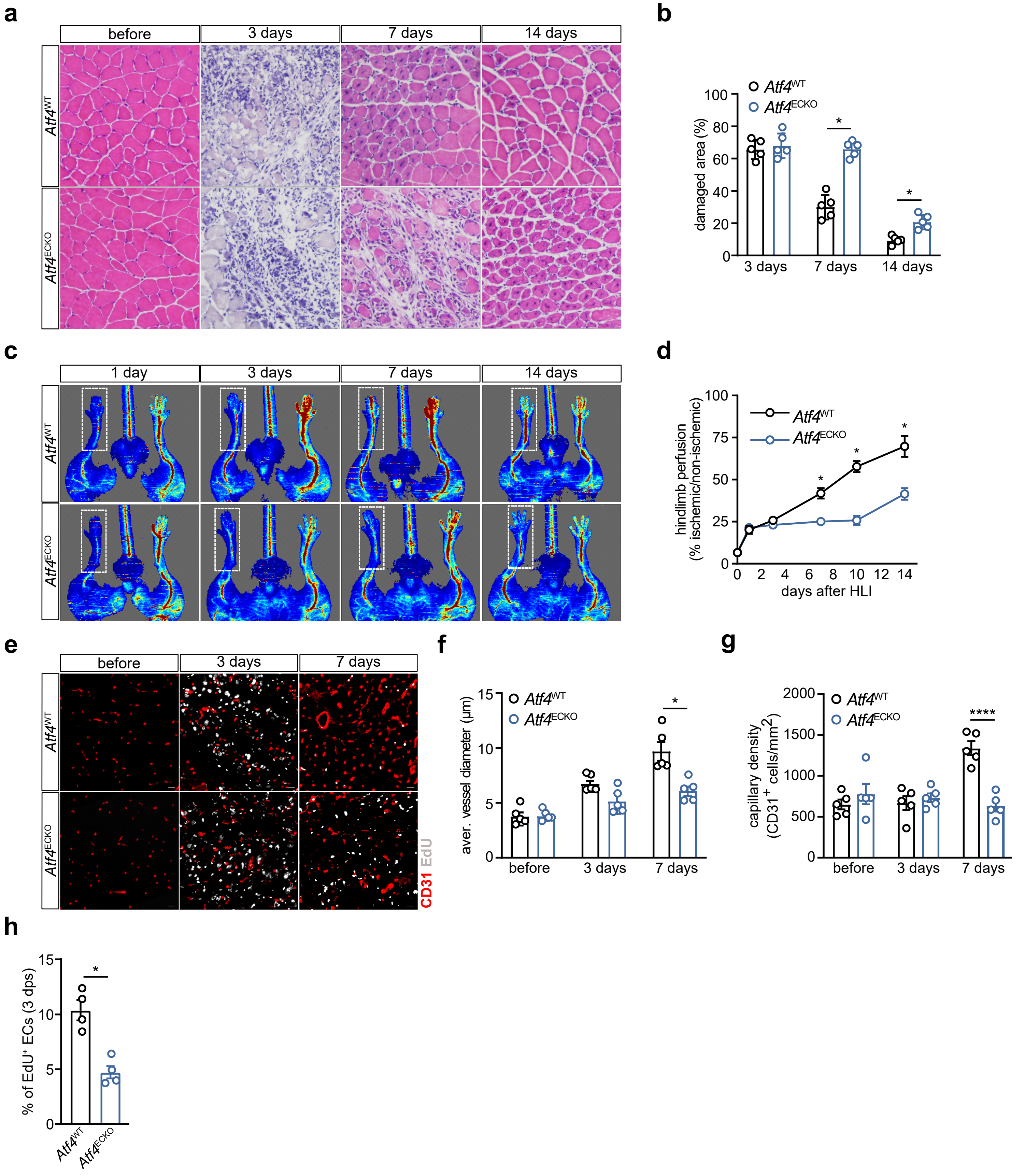

a
before
3 days
7 days
14 days
Atf4WT
Atf4ECKO
b
Atf4WT
Atf4ECKO
damaged area (%)
3 days
7 days
14 days
c
1 day
3 days
7 days
14 days
Atf4WT
Atf4ECKO
d
hindlimb perfusion (% ischemic/non-ischemic)
Atf4WT
Atf4ECKO
days after HLI
e
before
3 days
7 days
Atf4WT
Atf4ECKO
CD31 EdU
f
aver. vessel diameter (µm)
Atf4WT
Atf4ECKO
before
3 days
7 days
g
capillary density (CD31+ cells/mm2)
Atf4WT
Atf4ECKO
before
3 days
7 days
h
% of EdU+ ECs (3 dps)
Atf4WT
Atf4ECKO

**Extended Data Fig. 3: ATF4 is required in ECs for ischemia-induced revascularization in a preclinical model of PAD. a,** Representative images of haematoxylin and eosin (H&E)-stained sections of gastrocnemius muscle from *Atf4*$^{WT}$ (n=5) and *Atf4*$^{ECKO}$ (n=5) mice. **b,** Quantification of percentage of regenerative area from panel a measured as areas with mononuclear cells infiltration and fibers with centrally localized nuclei. **c-d,** Representative laser Doppler images (**c**) and quantification of hindlimb perfusion (**d**) in *Atf4*$^{WT}$ (n=6) and *Atf4*$^{ECKO}$ (n=6) mice. **e,** Representative images of gastrocnemius muscle from *Atf4*$^{WT}$ (n=5) and *Atf4*$^{ECKO}$ (n=5) mice stained for the EC marker (CD31, red) and EdU (white) (Scale bar: 20 µm). **f-h**, Quantification of vessel diameter (**f**), vessel density (**g**) and percentage of proliferative (EdU$^{+}$) ECs (**h**) from panel e. Each dot represents a single mouse in panels b, f, g and h or mean ± SEM in panel d. Two-way ANOVA with Sidak's multiple comparisons ($^{*}p < 0.05$) was used in b, d, f and g. Student's t test (two-tailed, unpaired, parametric, $^{*}p < 0.05$) was used in h. P = 0.0007 (**d**, 7 days), P < 0.0001 (**d**, 10 days), P < 0.0001 (**d**, 14 days), P < 0.0001 (**f**, 7 days), P < 0.0001 (**g**, 7 days), P = 0.0019 (**h**).

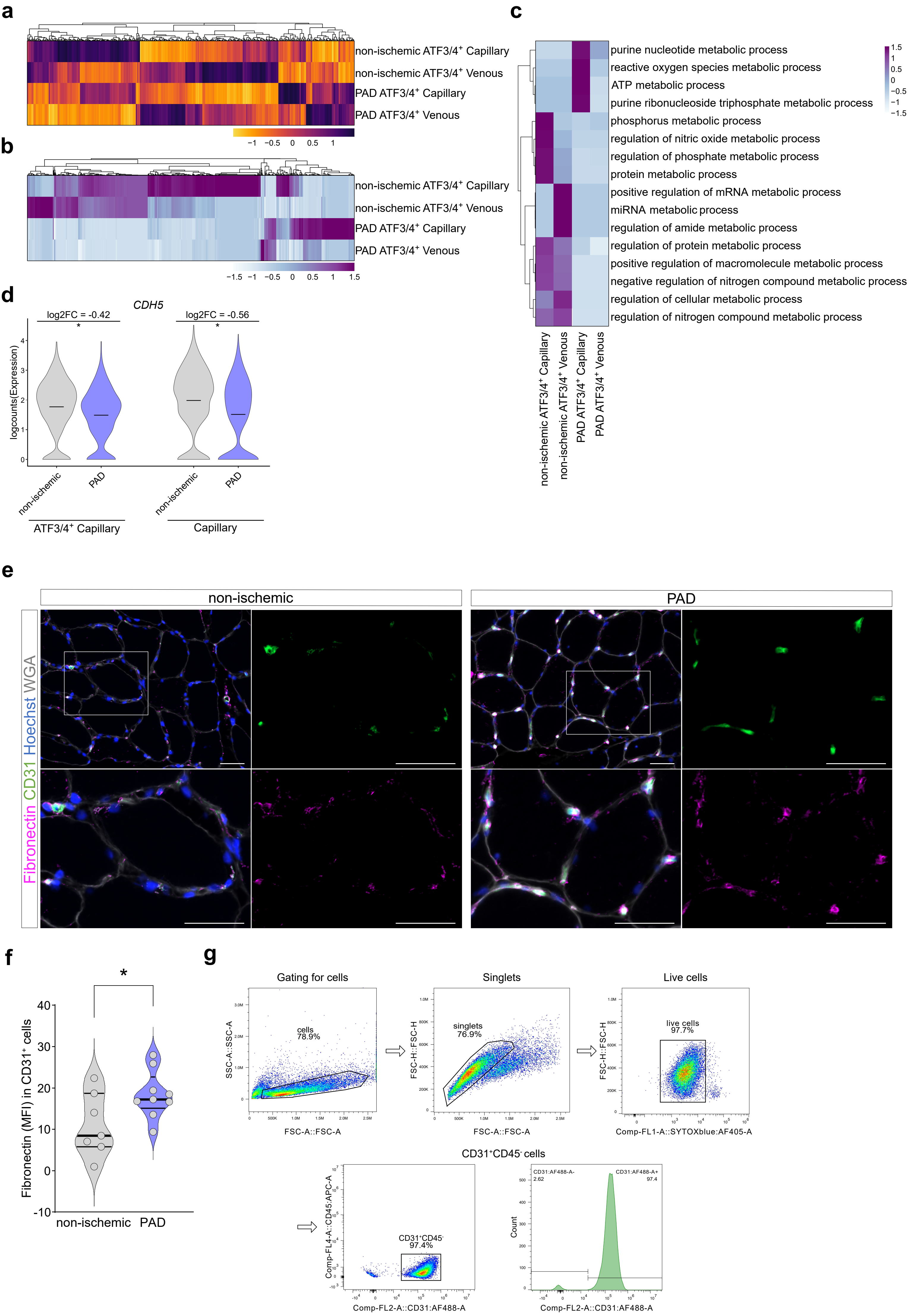
a
non-ischemic ATF3/4+ Capillary
non-ischemic ATF3/4+ Venous
PAD ATF3/4+ Capillary
PAD ATF3/4+ Venous
−1 −0.5 0 0.5 1
b
non-ischemic ATF3/4+ Capillary
non-ischemic ATF3/4+ Venous
PAD ATF3/4+ Capillary
PAD ATF3/4+ Venous
−1.5 −1 −0.5 0 0.5 1 1.5
c
purine nucleotide metabolic process
reactive oxygen species metabolic process
ATP metabolic process
purine ribonucleoside triphosphate metabolic process
phosphorus metabolic process
regulation of nitric oxide metabolic process
regulation of phosphate metabolic process
protein metabolic process
positive regulation of mRNA metabolic process
miRNA metabolic process
regulation of amide metabolic process
regulation of protein metabolic process
positive regulation of macromolecule metabolic process
negative regulation of nitrogen compound metabolic process
regulation of cellular metabolic process
regulation of nitrogen compound metabolic process
1.5 1 0.5 0 −0.5 −1 −1.5
non-ischemic ATF3/4+ Capillary
non-ischemic ATF3/4+ Venous
PAD ATF3/4+ Capillary
PAD ATF3/4+ Venous
d
CDH5
log2FC = -0.42
log2FC = -0.56
logcounts(Expression)
non-ischemic
PAD
ATF3/4+ Capillary
Capillary
e
non-ischemic
PAD
Fibronectin CD31 Hoechst WGA
f
Fibronectin (MFI) in CD31+ cells
non-ischemic PAD
g
Gating for cells
cells 78.9%
SSC-A::SSC-A
FSC-A::FSC-A
Singlets
singlets 76.9%
FSC-H::FSC-H
FSC-A::FSC-A
Live cells
live cells 97.7%
FSC-H::FSC-H
Comp-FL1-A::SYTOXblue:AF405-A
CD31+CD45- cells
CD31+CD45- 97.4%
Comp-FL4-A::CD45:APC-A
Comp-FL2-A::CD31:AF488-A
CD31:AF488-A- 2.62
CD31:AF488-A+ 97.4
Count
Comp-FL2-A::CD31:AF488-A

**Extended Data Fig. 4: *ATF3/4*$^+$ ECs undergo a profound transcriptomic rewiring in PAD. a**, Heatmap of centered logcounts of DEGs (adjusted p-value <0.05) between *ATF3/4*$^+$ ECs in non-ischemic and PAD, color indicates the centered logcount value. **b-c**, Heatmap of centered values from ORA analysis over the DEGs in *ATF3/4*$^+$ Venous and Capillary ECs between conditions, color indicates the centered values. Panel b displays all the enriched processes and panel c focuses on metabolic processes. **d**, Violin plots showing logcounts values of *CDH5* expression between conditions, a negative log2FC indicates a downregulation in PAD ($^*p < 0.05$), horizontal line indicates the average logcounts value. **e,** Representative images of gastrocnemius muscle from non-ischemic (n=7) and PAD (n=9) patients stained for Fibronectin (red), the EC marker (CD31, green) cell nuclei (Hoechst, blue) and WGA (white) (Scale bar: 50 µm). In each group, the top left panel has a lower magnification compared to the other 3 pictures. White rectangle indicates the zoomed area displayed in the remaining panels **f,** Quantification of Fibronectin mean fluorescence intensity (MFI) in CD31$^+$ cells from panel e. **g,** Flow cytometric analysis of percentage of CD31$^+$CD45$^-$ cells in primary cells isolated from human gastrocnemius muscle (related to Fig. 3, see Methods). Each dot represents a single patient in panel f. Wilcoxon Rank Sum test (as implemented in Seurat package) was used in a and d. Student's t test (two-tailed, unpaired, parametric, $^*p < 0.05$) was used in f. P = 1.04e-19 (**d**, *ATF3/4*$^+$ Capillary), P = 6.11e-39 (**d**, Capillary), P = 0.0454 (**f**).

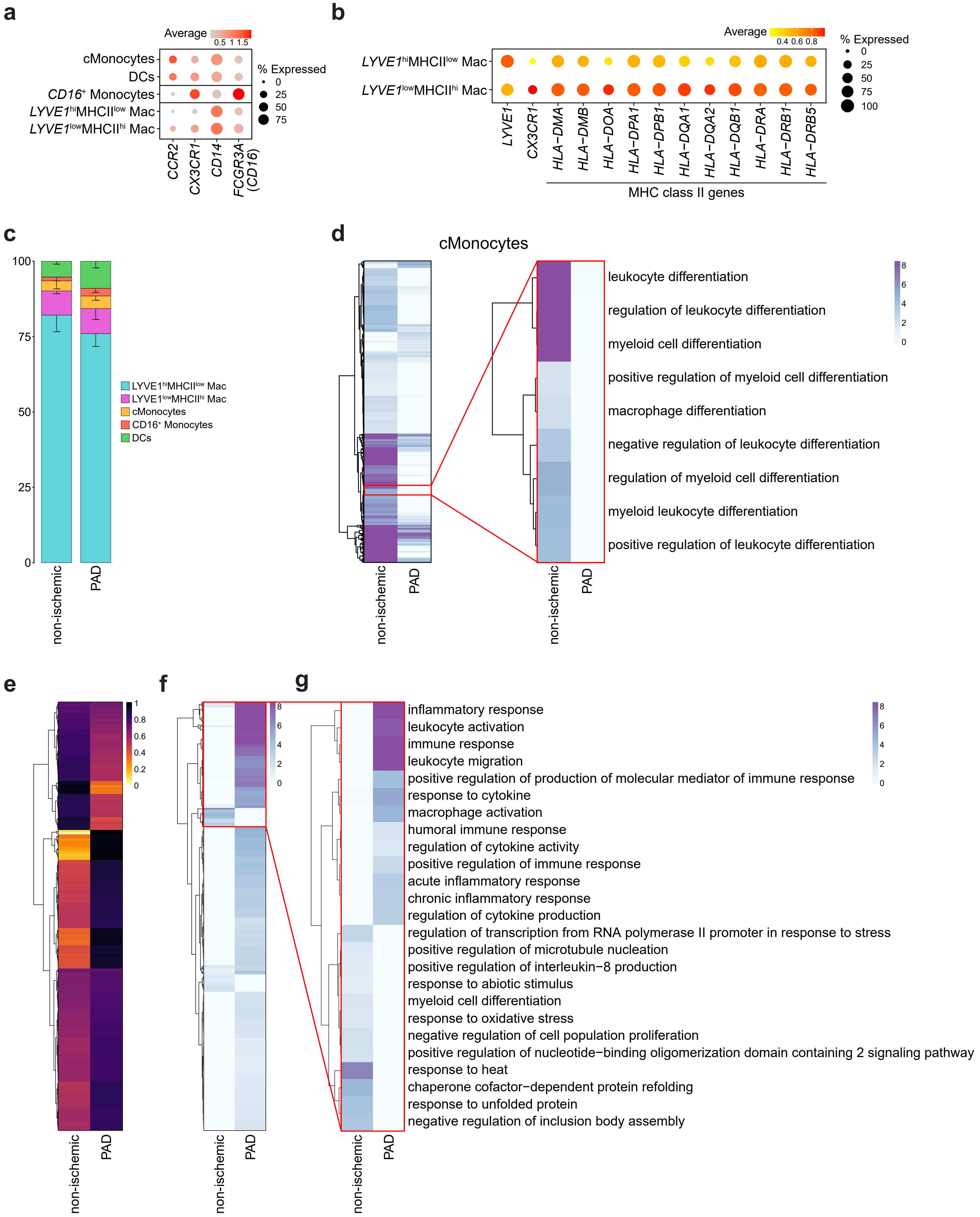
a
Average
0.5 1 1.5
% Expressed
0
25
50
75
cMonocytes
DCs
CD16⁺ Monocytes
LYVE1hiMHCIIlow Mac
LYVE1lowMHCIIhi Mac
CCR2
CX3CR1
CD14
FCGR3A (CD16)
b
Average
0.4 0.6 0.8
% Expressed
0
25
50
75
100
LYVE1hiMHCIIlow Mac
LYVE1lowMHCIIhi Mac
LYVE1
CX3CR1
HLA−DMA
HLA−DMB
HLA−DOA
HLA−DPA1
HLA−DPB1
HLA−DQA1
HLA−DQA2
HLA−DQB1
HLA−DRA
HLA−DRB1
HLA−DRB5
MHC class II genes
c
100
75
50
25
0
LYVE1hiMHCIIlow Mac
LYVE1lowMHCIIhi Mac
cMonocytes
CD16⁺ Monocytes
DCs
non-ischemic
PAD
d
cMonocytes
leukocyte differentiation
regulation of leukocyte differentiation
myeloid cell differentiation
positive regulation of myeloid cell differentiation
macrophage differentiation
negative regulation of leukocyte differentiation
regulation of myeloid cell differentiation
myeloid leukocyte differentiation
positive regulation of leukocyte differentiation
8
6
4
2
0
non-ischemic
PAD
e
1
0.8
0.6
0.4
0.2
0
non-ischemic
PAD
f
8
6
4
2
0
non-ischemic
PAD
g
inflammatory response
leukocyte activation
immune response
leukocyte migration
positive regulation of production of molecular mediator of immune response
response to cytokine
macrophage activation
humoral immune response
regulation of cytokine activity
positive regulation of immune response
acute inflammatory response
chronic inflammatory response
regulation of cytokine production
regulation of transcription from RNA polymerase II promoter in response to stress
positive regulation of microtubule nucleation
positive regulation of interleukin−8 production
response to abiotic stimulus
myeloid cell differentiation
response to oxidative stress
negative regulation of cell population proliferation
positive regulation of nucleotide−binding oligomerization domain containing 2 signaling pathway
response to heat
chaperone cofactor−dependent protein refolding
response to unfolded protein
negative regulation of inclusion body assembly
8
6
4
2
0
non-ischemic
PAD

**Extended Data Fig. 5: *LYVE1*$^{hi}$MHCII$^{low}$ macrophages get activated in PAD. a**, Dot plot of logcount values of markers from different myeloid populations in Monocytes/Macrophages subtypes. Color and size of the dots indicate the logcount value and the proportion of cells that express the gene, respectively. **b**, Dot plots of logcounts values of *LYVE1*, *CX3CR1* and MHC-II class genes in *LYVE1*$^{hi}$MHCII$^{low}$ and *LYVE1*$^{low}$MHCII$^{hi}$ macrophages. Color and size of the dots indicate the logcount value and the proportion of cells that express the gene, respectively. **c**, Stacked bar plots showing cluster percentage in each condition in the spatial dataset from non-ischemic (n=3) and PAD (n=3) patients, color-coded by cluster. Each stack represents mean - SEM. **d,** Heatmap of centered values from ORA analysis over the DEGs in cMonocytes between conditions, color indicates the centered values. The left heatmap shows all identified enriched processes, while the right heatmap only shows terms related to leukocyte/myeloid cell/macrophage differentiation. **e,** Heatmap of logcounts values of DEGs (adjusted p-value <0.05) between *LYVE1*$^{hi}$MHCII$^{low}$ macrophages in non-ischemic and PAD, color indicates the logcount value. **f-g,** Heatmap of centered values from ORA analysis over the DEGs in *LYVE1*$^{hi}$MHCII$^{low}$ macrophages between conditions, color indicates the centered values. Panel f displays all the enriched processes and panel g focuses on top enriched processes in each condition. Wilcoxon Rank Sum test (as implemented in Seurat package) was used in e.

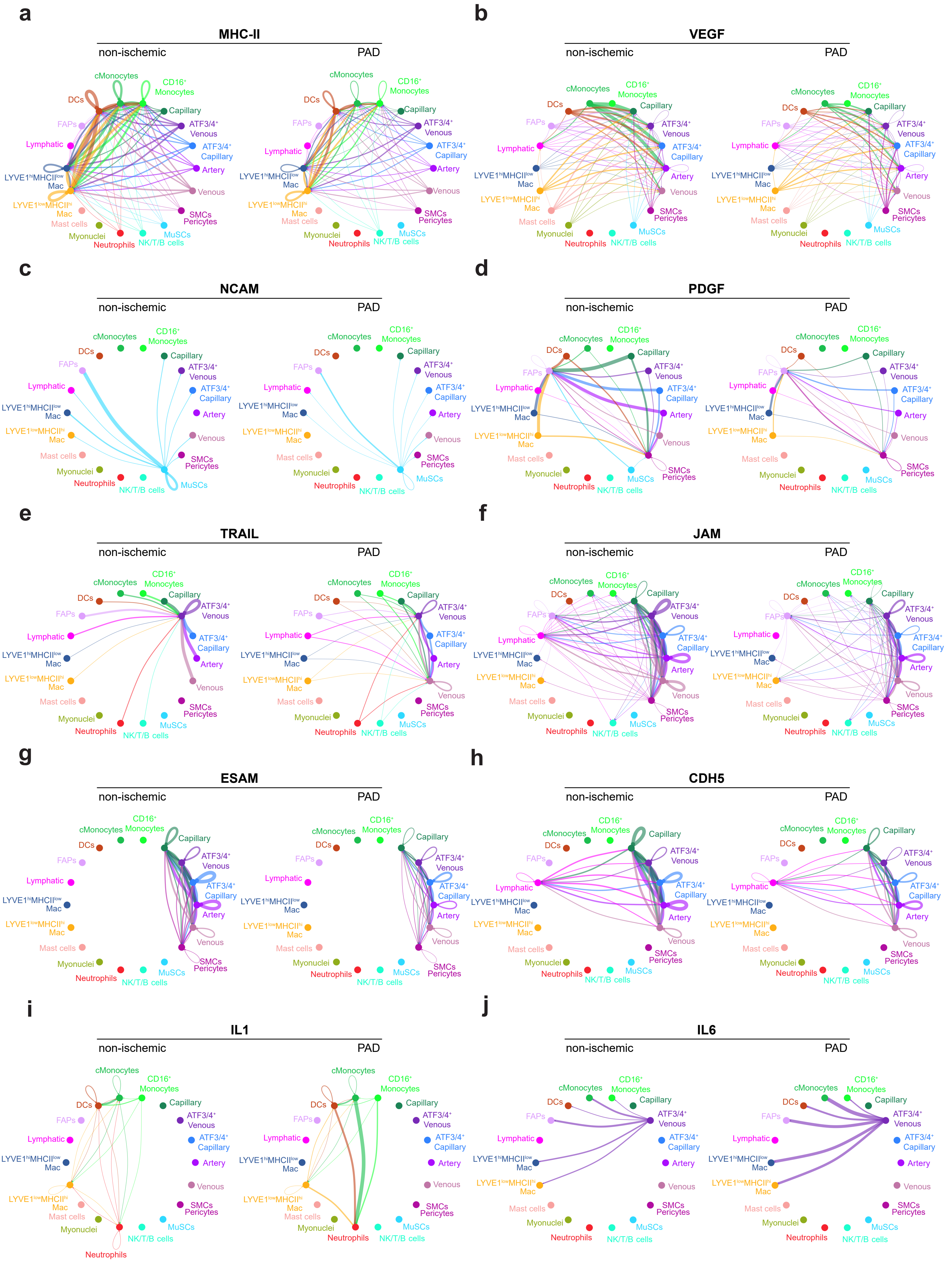

a
MHC-II
b
VEGF
c
NCAM
d
PDGF
e
TRAIL
f
JAM
g
ESAM
h
CDH5
i
IL1
j
IL6
non-ischemic
PAD
cMonocytes
CD16$^+$ Monocytes
DCs
Capillary
FAPs
ATF3/4$^+$ Venous
Lymphatic
ATF3/4$^+$ Capillary
Artery
LYVE1$^{hi}$MHCII$^{low}$ Mac
Venous
LYVE1$^{low}$MHCII$^{hi}$ Mac
SMCs Pericytes
Mast cells
Myonuclei
MuSCs
Neutrophils
NK/T/B cells

**Extended Data Fig. 6: Cellular communication in human skeletal muscle is disrupted in PAD. a-j,** Circle plots showing MHC-II (**a**), VEGF (**b**), NCAM (**c**), PDGF (**d**), TRAIL (**e**), JAM (**f**), ESAM (**g**), CDH5 (**h**), IL1 (**i**) and IL6 (**j**) communication in non-ischemic (left) and PAD (right). Color and width of the edges indicates the sender and weight of interactions, respectively.

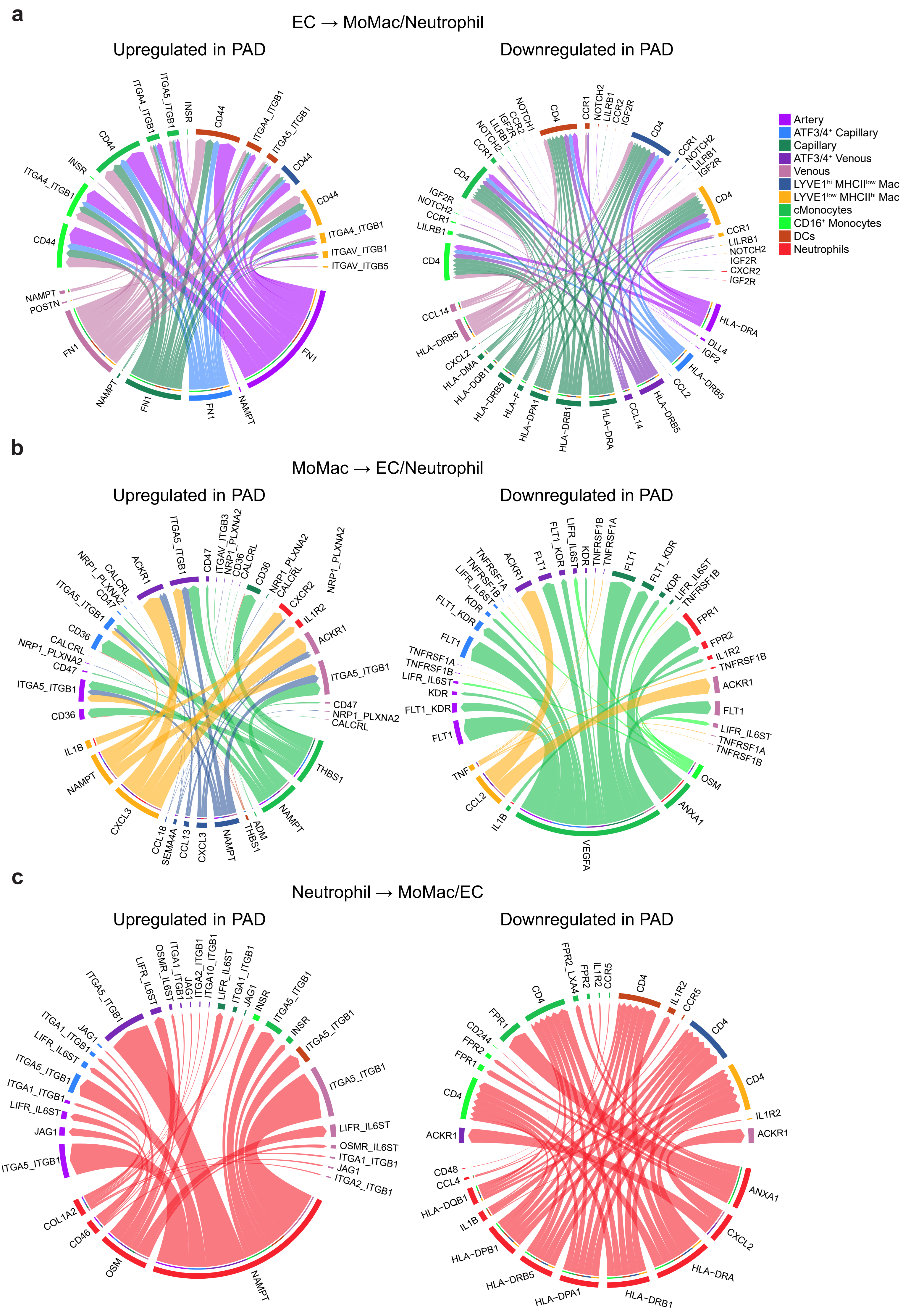
a
EC → MoMac/Neutrophil
Upregulated in PAD
Downregulated in PAD
b
MoMac → EC/Neutrophil
Upregulated in PAD
Downregulated in PAD
c
Neutrophil → MoMac/EC
Upregulated in PAD
Downregulated in PAD
Artery
ATF3/4⁺ Capillary
Capillary
ATF3/4⁺ Venous
Venous
LYVE1ʰⁱ MHCIIˡᵒʷ Mac
LYVE1ˡᵒʷ MHCIIʰⁱ Mac
cMonocytes
CD16⁺ Monocytes
DCs
Neutrophils
ITGA4_ITGB1
ITGA5_ITGB1
INSR
CD44
ITGAV_ITGB1
ITGAV_ITGB5
NAMPT
POSTN
FN1
NOTCH1
NOTCH2
CCR1
CCR2
LILRB1
IGF2R
CD4
CXCR2
HLA-DRA
DLL4
IGF2
HLA-DRB5
CCL2
CCL14
HLA-DRB1
HLA-DPA1
HLA-F
HLA-DQB1
HLA-DMA
CXCL2
ITGAV_ITGB3
NRP1_PLXNA2
CD36
CALCRL
CD47
ACKR1
CXCR2
IL1R2
THBS1
ADM
CXCL3
CCL13
SEMA4A
CCL18
IL1B
TNFRSF1A
TNFRSF1B
LIFR_IL6ST
KDR
FLT1_KDR
FLT1
FPR1
FPR2
OSM
ANXA1
VEGFA
TNF
OSMR_IL6ST
ITGA1_ITGB1
ITGA2_ITGB1
ITGA10_ITGB1
JAG1
COL1A2
CD46
FPR2_LXA4
CCR5
CD244
CD48
CCL4
HLA-DPB1

**Extended Data Fig. 7: Reciprocal communication between EC-MoMac-Neutrophil promotes pro-inflammatory and anti-angiogenic communication during PAD. a-c,** Circle plots showing upregulated (left panel) or downregulated (right panel) communication during PAD from ECs (**a**), MoMacs (**b**) and Neutrophils (**c**) to the other cell types. Color and width of the edges indicates the sender and weight of interactions, respectively.
